# Evolutionarily conserved waves of tooth replacement in the gecko are dependent on local signaling

**DOI:** 10.1101/2022.11.17.513312

**Authors:** Kirstin S. Brink, Eric Cytrynbaum, Theresa M. Grieco, Joaquin I. Henriquez, Anna Zhitnitsky, Joy M. Richman

**Affiliations:** Department of Oral Health Sciences, University of British Columbia, Canada; Department of Geological Sciences, University of Manitoba, Canada; STEMCELL Technologies, Vancouver, BC, CANADA; Department of Physics of Complex Systems, Weitzman Institute, Israel; Department of Mathematics, University of British Columbia, Canada

**Keywords:** reptile, dentition, polyphyodont, successional teeth, patterning mechanisms, mathematical modelling

## Abstract

The fossil record contains dinosaur jaws with rows of unerupted successional teeth that are arranged in a variety of elegant patterns. The remnants of these patterns are visible in modern dentate reptiles but the mechanism for generating and maintaining these rows of teeth is unknown. The biology underlying the tooth replacement pattern was hypothesized to either be stimuli transmitted across tooth families in the jaws (Edmund) or secretion of local inhibitory molecules that would stagger development of adjacent tooth families (Osborn). To test these hypotheses and generate new ones, we completed a study on 6 treated adult geckos in which one side of the jaw had teeth removed. Wax bites were used to record the maxillary teeth 2 times a week. Tooth presence or absence was recorded and transformed mathematically. The time between eruption at each tooth position was measured as was the relative phase compared to the immediate adjacent teeth over successive bites. The period between eruption events at each tooth position was approximately 30 days with some lengthening over time. The average relative phase showed there was a tilt in the data that fit the observation that alternating teeth were being shed. This tilt was opposite on the left and right sides of the jaw. The asymmetry of the right and left sides was consistent across the dentition. After plucking, the pattern recovers after 3 periods fitting with the consistent finding that there are 3 teeth in each tooth family. Ablated areas did not recover tooth formation even after 14 months. The plucked animals showed evidence of fixed, local signaling that restores the pattern. Two models based on Osborn’s concept of a “zone of inhibition” deviate from the observed data. The ablated animals show no change in patterns of tooth eruption anterior and posterior to the gap. Thus there is no support for the Wave stimulus theory of Edmund. Finally, we propose a new Phase Inhibition Model. This model assumes fixed initiation sites at which teeth are initiated at some phase within a month-long cycle and that, as a tooth is initiated, the cycles of nearby initiation sites are inhibited in their progress. This inhibition causes nearest neighbours to erupt in anti-synchrony. This model best maintained the tilt, spacing timing of the real biological data. Mathematical modeling was sensitive enough to measure the normal developmental instability and the resilience of the gecko to restore homeostasis after tooth removal.

## Introduction

The principals of generating repeated biological patterns have been derived from careful observations of developing organisms from *Drosophila*, frog, chicken and mouse. Mathematical scientists starting in the 20^th^ century have thought deeply about biological patterning based on observations of nature. Allan Turing was one of the most influential mathematicians to postulate a mechanism that could explain repeated patterns (Economou and Green 2014). Turing suggested that patterns could be set up with just two chemicals - an activator and a faster diffusing inhibitor thus creating the reaction-diffusion model (Turing 1952). The model predicted the formation of standing waves. The peaks and valleys determined where the structures were formed and the space between them. The model presumes that the level of each chemical is stable and thus once formed the pattern would also be maintained. However, reaction-diffusion may not apply to all biological patterns. For example, how does reaction-diffusion operate in organs such as hair that have distinct spacing and polarity and yet undergo dynamic renewal in the adult animal (Aw and Devenport 2017). Furthermore, from what is known about the properties of growth factors, long range diffusion is less likely to be involved in adult organisms with highly differentiated structural matrices. There is a need for more diverse types of animal data that can be used to test models of patterning.

In this study we focus patterning in the adult dentition. Although teeth are only replaced twice in humans and other mammals, the reptiles have numerous taxa that exhibit polyphyodonty (tooth replacement throughout life). Reptiles are amniotes and thus share a common ancestor with mammals(Reisz 1997). Therefore, ontogeny of teeth in squamate reptiles and crocodilians is conserved with mammals, following the typical stages from initiation to bud, cap and bell stages (Buchtová et al. 2007; Buchtová et al. 2008; Buchtova et al. 2013; Jernvall and Thesleff 2012; Wu et al. 2013). The ‘waves’ of tooth replacement in adult animals have been described by others for more than a century from skulls (Edmund 1960; 1962; Woerdeman 1919). However, there are only rare studies that follow tooth replacement in living reptiles. The adult iguana and alligator were followed using monthly radiographs for more than 2 years (Brink et al. 2020; Edmund 1962). Several lizards have been tracked using wax bites (Cooper 1966; Kline and Cullum 1984; Kline and Cullum 1985; Osborn 1971). More recently microCT was used to track mineralized tooth replacement in 3 alligators, 3 times in 1 year (Widelitz et al. 2017). The results in iguanas (Kline and Cullum 1984; Kline and Cullum 1985), anguis fragilis (slow worm) (Cooper 1966) and alligator (Edmund 1962) were similar in that there were waves of replacement stretching from the posterior to the anterior of the jaw. Approximately every other tooth was shed, leaving some teeth to function at all times. The variability of the patterns of shedding appeared to increase as the animal aged. Osborne was the person to publish most extensively on the mathematical description of tooth replacement estimated that in the iguana and alligator it took about 3-4 months for a replacement tooth to progress from imitation to function (Osborn 1974). However, he was limited by the number of teeth in the jaw of these species and their very slow rate of replacement.

Mathematical modeling of tooth replacement has only been carried out in a limited number of studies. The Edmund model proposed a source of patterning information that diffuses from the anterior to posterior of the jaw or the wave-stimulus model (Edmund 1960; 1962; Whitlock and Richman 2013). Edmund did not generate equations to represent his observations and inserted lines connecting the teeth that could have been drawn in many different ways. The Osborn model (Osborn 1974) examined lizard data (Cooper 1966) as well as cross-sectional data from skeletal remains of different aged animals from Edmund (Edmund 1960). Osborne wondered how the stable dynamic shape of the ‘wave of tooth replacement of alternate teeth’ was maintained throughout life? He did not consider unerupted teeth but only those present in the mouth. Osborn tested the possibility that adjacent teeth were lost sequentially but found that the biology did not fit this mathematical model. He also ruled out a model where adjacent teeth were at exactly the same stage of development. For his final model he proposed a zone of inhibition radiating out from the developing teeth. The radius of this inhibitory zone might be the reason for adjacent successional teeth being at different stages of development in biological observations (Edmund 1960). This would lead to a group of neighbouring teeth being shed at the same time which was never observed in the jaws. Osborn proposed that the inhibitory zones are repeated in every tooth family and this would organize posterior-to-anterior waves of tooth replacement. He also thought that over time, the size of teeth in successive generations would increase because they would sit in a stage of initiation for longer. The hypotheses of Osborn need to be tested rigorously across a group of animals of the same species that has captured tooth shedding over a sufficient period of time in a jaw with many tooth positions.

We have developed the leopard gecko as our model to study adult tooth replacement because this animal has advantages over other reptiles. First, the animals are relatively easy to raise in a laboratory setting (Vickaryous and Gilbert 2019), they are amenable to taking wax bites, a non-invasive way to follow in a consistent pattern and all teeth have a simple morphology (homodont)(Grieco and Richman 2018; Handrigan and Richman 2011). The replacement rate in geckos is much more rapid (4-5 weeks) than iguana(Brink et al. 2020) or alligator(Edmund 1962). There are 40 teeth in each quadrant of the gecko rather than 17-20 in alligators and iguanas thus the density of tooth shedding events is higher (Brink et al. 2020; Edmund 1962). These additional data points will increase sensitivity to detect jaw-wide or local alterations in the patterning of tooth shedding events.

Only two of the reptile tooth replacement studies from the past included experiments that altered tooth number and followed the effects over time. The first study was carried out by Edmund on iguanas but not analyzed until our group re-examined the original radiographs (Brink et al. 2020). Edmund had removed functional, erupted teeth in several iguanas and studied healing thereafter. There was no effect on the rate of tooth replacement (20 weeks between shedding events for a single tooth). The study by Cooper (Cooper 1966) did not experimentally perturb the iguanas but happened to note there was a tooth loss event during the study that self-corrected. One study selectively extracted functional teeth in juvenile alligators (Wu et al. 2013) but the animals were euthanized 4 weeks after removal of erupted teeth. Thus the long term effects on tooth succession were not explored.

The leopard gecko is a good model in which to carry out experimental perturbations. We have developed methods to selectively remove a set of unerupted teeth to offset the timing of tooth initiation, eruption and shedding in the centre of the jaw (Brink et al. 2021). We previously showed that animals recover the normal dentition within 3 months of removing the unerupted, second generation teeth. However, in this previous study (Brink et al. 2021) we had not followed the animals for longer than 3 months. Instead attention was focused on the cellular events that occurred after tooth removal and the contribution of epithelial stem cells to the next generation of teeth (Brink et al. 2021). In the present study we combine the selective tooth removal experiments with long term tracking of tooth shedding. The aim is to test whether there are primarily local or long distance signaling that maintains the timing of tooth shedding. By removing teeth in the centre of the jaws we can test whether there is disruption of shedding anterior or posterior to the defect. In addition, we will determine how the gap is healed, whether from the edges or the centre. The type of healing will provide insights about the location of putative stem cells.

In order to gain an unbiased view of all the tooth shedding events over a long period of time we gathered biweekly wax bite data on multiple leopard geckos (approximately 4000 data points each for 6 animals). The raw scores were transform to mathematical representations of the biological data. We then went further to experimentally change the number of successor teeth by plucking (enucleated) or permanently preventing regrowth using chemical cauterization of the dental epithelium. We tested compared the mathematically transformed data to the predictions made by the Edmond and Osborn models. We developed five alternative models of tooth development and tested the viability of each model as a mechanistic explanation for the observed spatiotemporal patterns. Ultimately the newly described ‘phase-inhibition’ model best fit the biological observations.

## Methods

The leopard geckos (*Eublepharis macularius*) were purchased as eggs from Just Geckos, Rocklin, CA and raised at the animal unit at the University of British Columbia. All experimental work is approved under the animal ethics protocol #A19-0308. Animals ranged in age from 1-3 years post-hatching (Table 1). The methods of removing plucking out or enucleating second generation teeth are as published (Brink et al. 2021)(Fig. 1A-G). Geckos recover the cycle of tooth replacement after a period of disruption that lasts approximately 3 months (Brink et al. 2021). Long term block in tooth replacement was produced by chemical cautery of the dental lamina using topical application of 15% Fe_2_(SO_4_)_3_ liquid to the area. (Fig. 1H). Wax bites were taken twice a week using pink base plate wax as published (Grieco and Richman 2018) (#H00812, Coltene, Whaledent, Inc., OH)(Fig. 1I). To highlight the indents, the wax bite was stained with Toluidine blue (Fig. 1I). Each indent was scored as a tooth present (white boxes) or absent (black boxes) and plotted on excel sheets (Fig. 1J-L). The data from the spreadsheets was converted to the times between eruptions at the same tooth position (Fig. 1L, blue dots)..The short term effects were followed using histology and cell proliferation analysis at 1,2, 4, 8, 12, 36 and 56 weeks post treatment (Table 1). Methods used included staining for proliferating cells with either BrdU or PCNA and staining for the dental epithelium with antibodies to PITX2 as published (Brink et al. 2021).

**Table 1.** specimens used in this study.

| Pluckers |  |  |  |  |  |
| --- | --- | --- | --- | --- | --- |
| Animal number | Study ID | Timepoint after treatment | weeks post treatment | Use | BrdU? |
| LG190 | Plucker A | 7 months | 28 | μCT+histology | no |
| LG169 | Plucker B | 7 months | 28 | adopted |  |
| LG170 | Plucker C | 7 months | 28 | adopted |  |
| Ablators Fe<sub>2</sub>(SO<sub>4</sub>)<sub>3</sub> |  |  |  |  |  |
| LG 200 | Ablator A | 1 week | 1 | μCT+histology | No |
| LG 253 | Ablator B | 1 week | 1 | μCT+histology | No |
| **LG 190\*** | Ablator C<br>(also Plucker A) | 2 week | 2 | μCT+histology | No |
| LG 255 | Ablator D | 2 week | 2 | μCT+histology | No |
| LG 230 | Ablator E | 1 month | 4 | μCT+histology | Yes |
| LG 231 | Ablator F | 1 month | 4 | μCT+histology | Yes 1 month chase before euthanasia |
| LG 216 | Ablator G | 2 months | 8 | μCT+histology | Yes, 2 month chase before euthanasia |
| LG 224 | Ablator H | 3 months | 12 | μCT+histology | Yes |
| LG 225 | Ablator I | 3 months | 12 | μCT+histology | Yes |
| LG 174 | Ablator J | 7 months | 28 | adopted | No |
| LG 219 | Ablator K | 9 months | 36 | wax bites, μCT+histology | Yes 1 month chase before euthanasia |
| LG 178 | Ablator L | 14 months | 56 | wax bites, PTA contrast enhanced microCT | no |
| LG 148 | Ablator M | 14 months | 56 | wax bites, μCT+histology | Yes with 1 month chase before euthanasia |

\*LG190 was used for plucking and followed for 7 months. Then the same animal was treated with Fe<sub>2</sub>(SO<sub>4</sub>)<sub>3</sub> and used for molecular analysis at 2 weeks. Hence this animal is both Plucker A and Ablator C.

**Figure 1.**
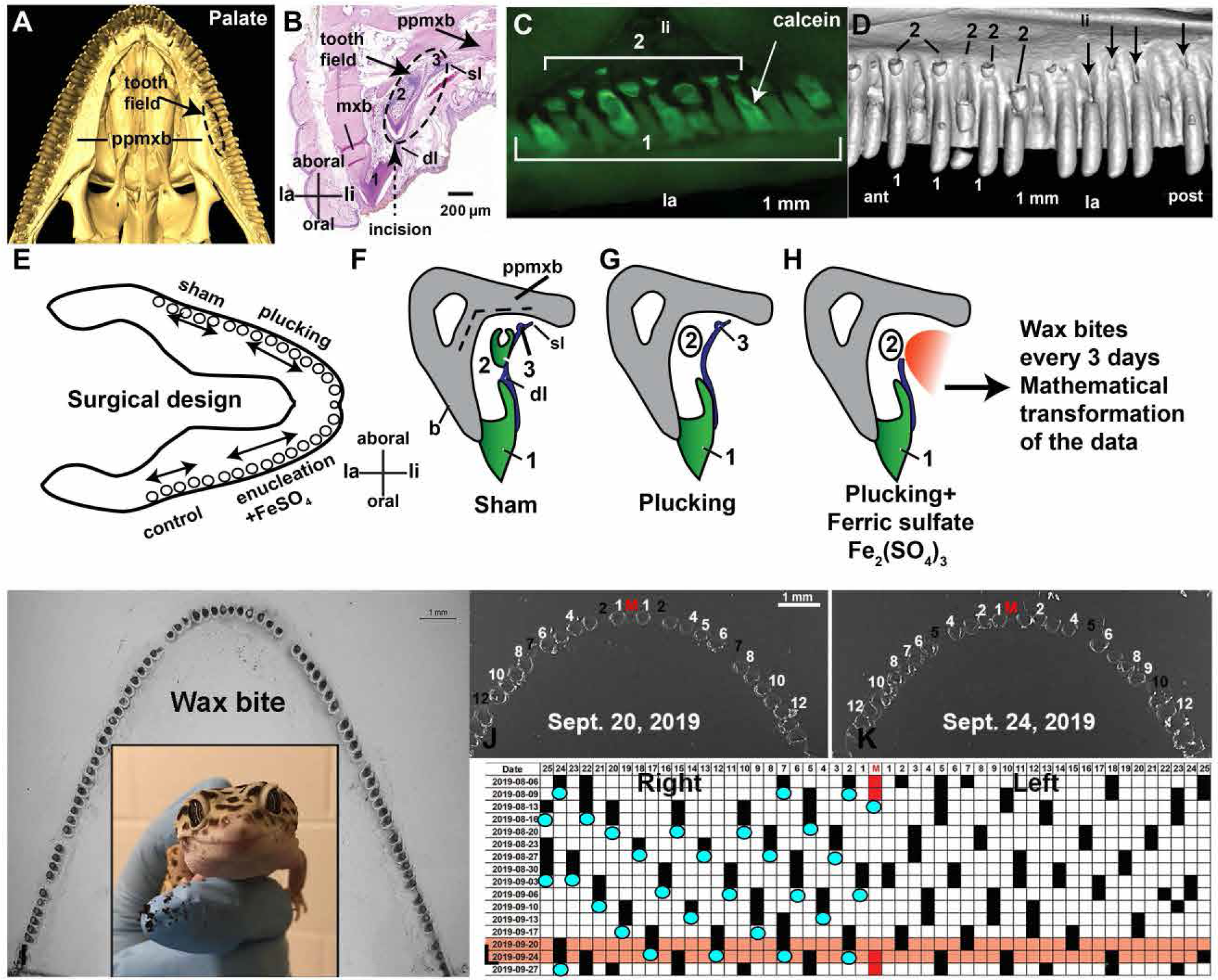
The experimental design for gecko surgeries. **A)** A μCT scan of a gecko palate showing the position of the tooth forming field (dashed line) relative to the functional teeth at the border of the maxillary bone. B) representative transverse section through the maxilla showing the outline of the tooth field (dashed line). C) Calcein is taken up by mineralizing dentin and enamel. Immature second generation teeth are visible under UV illumination. D) a μCT scan of an animal with teeth that were plucked (arrows). Adjacent non-treated area has 2^nd^ generation teeth (2). E) Scheme for dividing up the treatments into different parts of the jaw. For long term wax bites, only one treatment was carried out per animal, either plucking or Fe_2_(SO_4_)_3_ treatment. F-G) schematics of how plucking leaves the dental lamina behind and 3^rd^ generation teeth whereas Fe_2_(SO_4_)_3_ treatment eliminates most of the dental lamina. I) A wax bite stained with toluidine blue to highlight tooth indents. Gaps are where teeth are missing. J,K) Wax bites taken within the same week show differences in which teeth are present or absent. Not there is a single midline tooth in geckos (red M). L) Pink rows represent data in J, K. Blue spots are how data was transformed to represent eruption period. Key: 1 – first generation tooth in function, 2 – second generation tooth in late bell stage, 3 – third generation tooth in bud stage, ant-anterior, dl – dental lamina, la – labial, li – lingual, M – midline tooth, mxb – maxillary bone, post – posterior, ppmxb – palatine process of the maxillary bone, sl – successional lamina.

## Data representation

### Period heatmap

The time between tooth eruptions at each given tooth location (Fig. 1L) can be used to generate a period heat map, which gives information about the dynamic nature of tooth eruption periodicity across the jaw and in time.

### Phase heatmap

The phase of tooth eruptions can be represented much in the same way. It is calculated by taking the time of eruption of a given tooth as a ratio of the period of its neighbour. The phase can be calculated using left-neighbours or right neighbours, both of which conceptually convey the same information. We average the two directions to produce a measure of the regularity of the lattice, thereby removing information about the tilt (Fig. 2). Any regular checkerboard lattice (square or tilted) will give a uniform average phase of 0.5 period.

**Figure 2.**
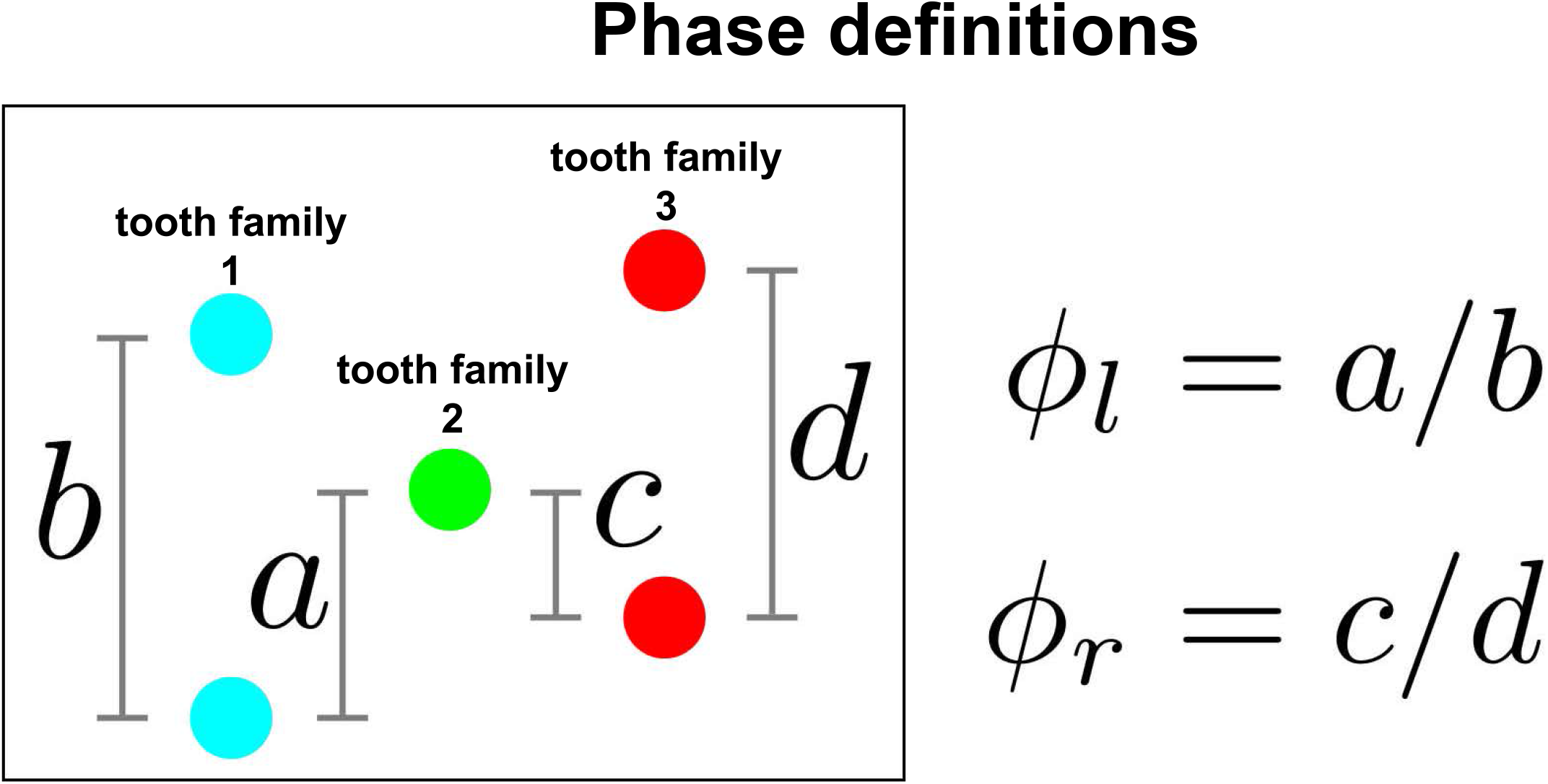
Calculating relative offset between neighbouring teeth. Three tooth families are illustrated with the more advanced teeth on top and the developing teeth at the bottom. The green tooth in tooth family 2 is used as the reference point. Measurements are made in between each member of the adjacent tooth families (blue and red). The spots in tooth families 1 and 3 do not align horizontally because they erupted at different times reflecting the real biological data. This calculation is repeated for all tooth positions to capture spatial and temporal data.

### Asymmetry heatmap

We observe a lattice tilt, which is mirrored about the centre of the jaw. To quantify this tilt, each tooth eruption is characterized by an asymmetry index which is measured by the difference in phase of a tooth relative to its left and right neighbour. These three representations of the data each highlight a different feature of interest and can be used to more closely analyze the validity of models. In particular, given the very noisy nature of these data, such heatmaps provide a way of making an otherwise qualitative analysis more robust.

## Model implementation

### Osborn’s Zone of Inhibition

The locations of new teeth in Osborn’s model are informed by the intersection of (inhibition) circles surrounding teeth of the preceding generation, because this is the earliest possible location not within the circles of inhibition; a lattice pattern of teeth thus emerges (Fig. 3). The results are then scaled by some constant velocity, *v*, at which the teeth move through the lamina. There are several factors that influence the patterns seen in this model, and are taken as parameters in the model. These are, *θ, d*, and *R*, which describe the angle between teeth, the distance between teeth and the radius of inhibition, respectively (Fig. 4). The tooth families (lines with short dashes) are slightly tilted (*θ* in Fig. 4) relative to vertical. The right and left side *θ* are antisymmetric. The zahnreihen connect every other tooth position, reflecting the tooth replacement patterns observed in many reptiles. Based on this, when we consider a jaw of finite size, we see periodic oscillatory patterns, where disturbances appear to reflect off the edges of the jaw due to the lack of inhibition there. At the edges of the jaw, the teeth are considered to be negatively interacting with the boundary; that is, the edge of the jaw is considered in the same way as the edge of an inhibiting circle.

**Figure 3.**
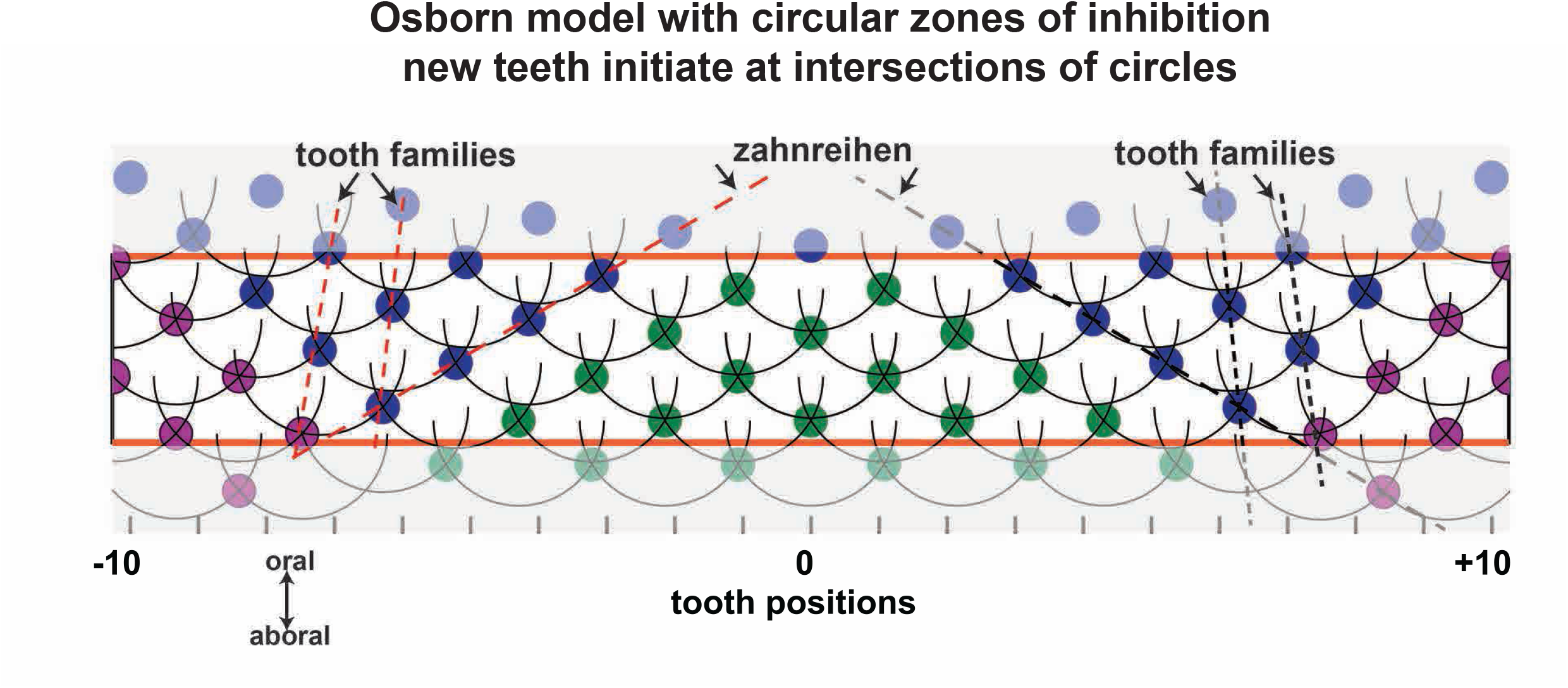
Osborn’s model with zones of inhibition. The locations of new teeth in Osborn’s model are informed by the intersection of (inhibition) circles surrounding teeth of the preceding generation because this is the earliest possible location not within the circles of inhibition. The tooth forming field is at the bottom of the illustration. There is no predetermined location for the initiating teeth. The top orange line indicates the oral mucosa where teeth will appear in the mouth. A lattice spacing of teeth emerges that steadily moves as new teeth are added. The results are scaled by a constant velocity, *v*, at which the teeth move through the lamina. Dashed line shows slight offset in the tooth family instead of being a straight vertical line. The diagonal lines show slightly offset successor teeth and the zahnreihen or waves of replacement connecting every other tooth position. Note that the right and left sides are mirror images of each other.

**Figure 4.**
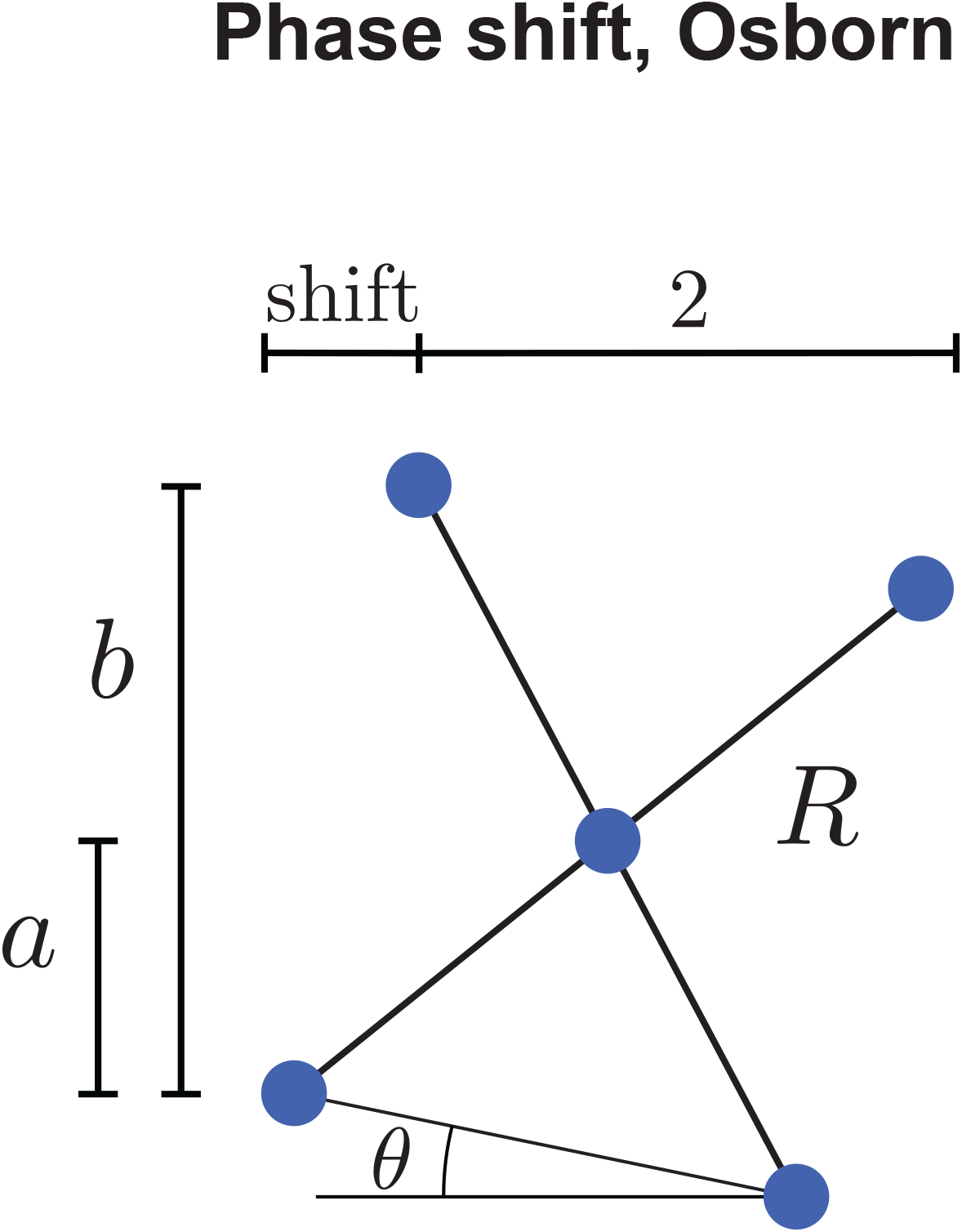
Osborn’s tilt. In the classic Osborn model, the teeth in tooth families cannot be in a vertical line due to the shape of the circles of inhibition. The geometry dictates that there is a slight tilt. This diagram shows 3 tooth families but unlike figure 2, the successor tooth is tilted by angle θ.

To simulate the plucker surgery, first a three-generation re-growth period is assumed based on the observation that there are typically three teeth lined up for each tooth location in zahnreihen. Following this delay period, teeth (and their inhibition circles) are removed (set to zero), so that teeth are allowed to re-grow in every integer tooth location in the surgical region. For the ablator surgery, the boundary of the surgical site is considered in the same way as the boundary of the jaw; the computation is effectively split into two parts.

### Phase & Phase Inhibition Model

The Phase and Phase Inhibition models are computed in the same way; in the former case the interaction parameter is simply set to zero. In addition, in order to initiate the computation, the phase model needs correct (stable) initial conditions in order to maintain a stable lattice, whereas the inhibition model converges to a stable configuration regardless of initial conditions. This is because the phase model does not have any means for converging or establishing a pattern which is needed in recovery from perturbations. For the plucker surgery, a three-generation recovery period is once again assumed based on biological data, after which teeth are allowed to regrow. Because this model computes in phase space, there are several possibilities for how teeth are allowed to regrow in the surgical region: a) the phase is unaffected (in which case the effect of the surgery is simply a gap in space and time; b) the phase is reset to zero to imitate a perfect initial condition and c) the phase is reset randomly. For the purpose of exploring how the model responds to perturbations, the phase of teeth following surgery is randomly reset. The ablator surgery is simulated in the same way as in the Osborn model; edges of the surgical site simply behave as boundaries in the same way as the edges of the jaw.

## Results

### Mathematical transformation of the plucker data

In order to analyze the entire dentition of multiple animals it was first necessary to transform the raw scores into a representation that captured the time it takes for a new tooth to appear/erupt at each location, the relative delays between adjacent teeth as well as the X and Y axis information (tooth positions and dates of bites). We imposed a heat map over each tooth position that varied in colour according to the interval between replacement of that tooth. The darker blue, the shorter the interval, the lighter blue colour indicates a slowing of the replacement rate (Plucker A, Fig. 5A).

**Figure 5.**
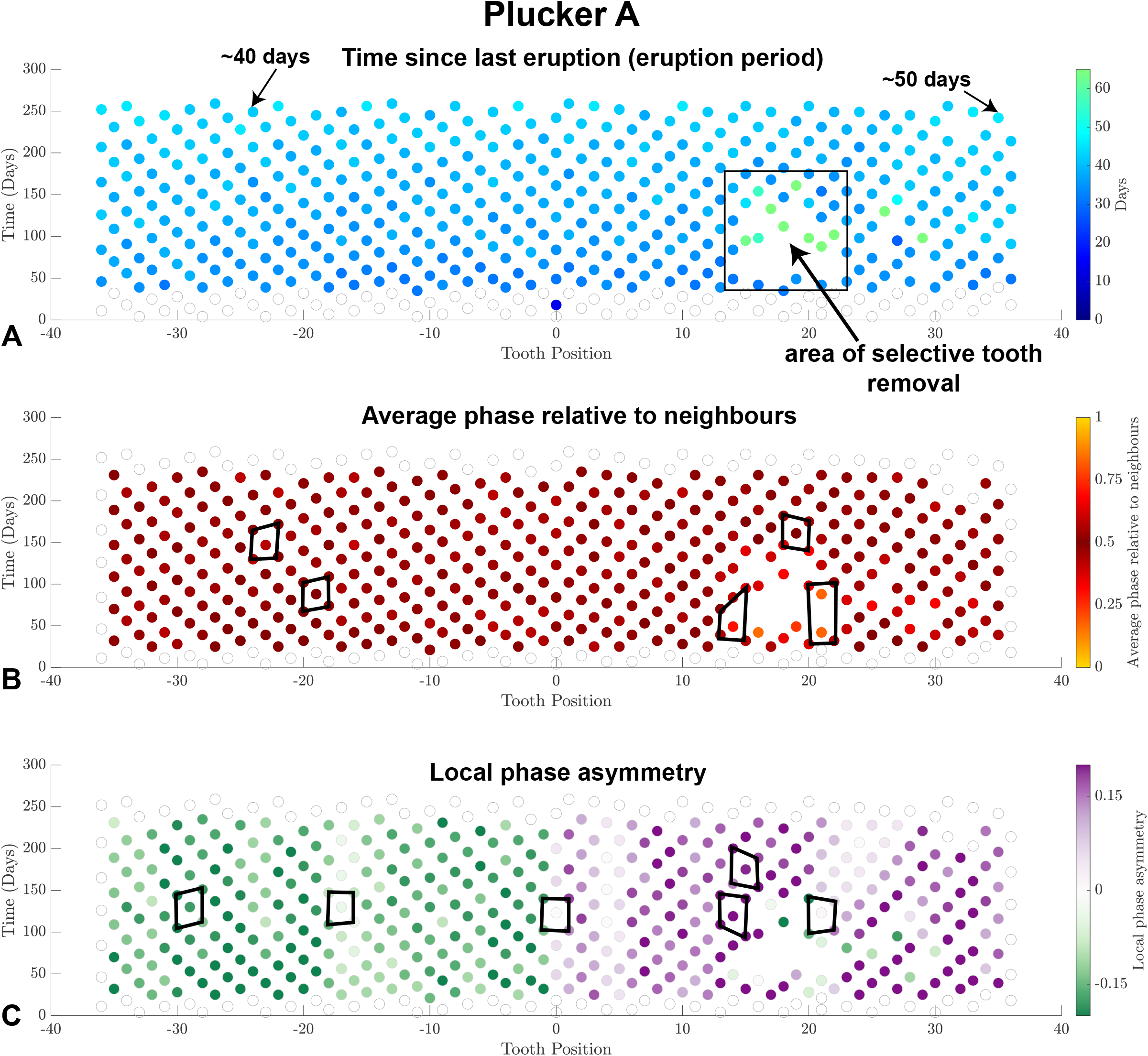
Plucker A showing the full post-surgery recovery period. A) The time between eruptions of teeth in a tooth family. Note teeth in a tooth family are slightly offset as in Figure 3. The heatmap shows length of time between eruptions. White circles are plotted where previous tooth information is lacking. B) Relative phase calculated between neighboring tooth families as in Figure 2. In all cases on the left side of the plot the tooth in the centre of the polygon is dark red reflecting a perfect half a phase offset. After the surgery the polygons are not as regular when connecting every other tooth position. Tooth in the centre is out of phase compared to the neighbours (more yellow). The position of the centre spot is also offset. The centre tooth erupts much closer in time relative to the immediate neighbours (closer together). C) Local phase asymmetry illustrates the mirror image offset on the right and left sides of the mouth. The offsets of the polygons reflect antisymmetry. The centre of the jaw is clearly identified (position 0) shows perfect squares that overlay nearly white teeth. Generally, the surgery temporarily disrupted phase asymmetry but this returned at 150 days or about 90 days since the surgery was carried out.

The representation of the biological data began by plotting a dot at the time since the previous eruption at the same location which we refer to as the eruption period (Fig. 1L). The eruption events are depicted with the horizontal component representing tooth position, measured as tooth number from the center, and vertical component representing the eruption time in days from the start of measurement (Fig. 5A). The last few tooth locations on both ends of the jaw were omitted from the plots because the wax bites were inconsistent in capturing those teeth. The colour of the dots in each panel indicates the value of each of the three metrics that we chose to characterize the patterns.

Focusing on the non-treated left side of the plot, there is a gradual increase in the eruption period over the span of months and possibly a slight increase in period going from the midline posteriorly along the jaw. To verify this, we calculated the best fit for the statistical model *E*(*N, t*) = *E*0 + *α*|*N* | + *βt* where *E*(*N, t*) is the eruption period at tooth location *N* and day *t*. For Plucker A, the left side of the jaw, we found a best fit for *E*0 = 30.1 (CI [29.3, 31.0]) days, *α* = 0.076 (CI [0.05, 0.10]) additional days per tooth location away from the center tooth and *β* = 0.046 (CI [0.041, 0.050]) additional days per day into the experiment. This means the center tooth had an eruption period of approximately 30.1 days initially rising to 41.6 days at day 250 of the experiment (30.1 + 0*α* + 250*β*) and the 36th tooth from the center had an eruption period of approximately 32.8 days initially (30.1 + 36*α* + 0*β*) rising to 44.3 days at day 250 of the experiment (30.1 + 36*α* + 250*β*). Other individuals showed evidence of similar patterns but not quite clearly as Plucker A (Figs. S1,S2). It is possible that the lack of clarityin the pattern is due to technical issues with the scoring.

Again focusing on the left, control side, the spatiotemporal eruption patterns also have a checkerboard-like nature to them with neighbouring teeth erupting asynchronously. In keeping with this checkerboard effect, there are apparent diagonal lines rising up and outward from the centerline of the jaw (Fig. 5A). The mirror-image up-and-inward lines that would normally appear on a checkerboard are also visible but are harder to detect because the inter-dot spacing is larger in the inward direction. To characterize the asynchrony and its asymmetry, we defined two quantities: average relative phase and asymmetry of relative phase. The left relative phase of an eruption is the ratio *ϕl* = *a/b* where *a* and *b* are defined (Fig. 2). The right relative phase is defined similarly, *ϕr* = *c/d*. The *average relative phase*, shown for the Plucker A eruption data (Fig. 5B) is the average of the left and right relative phases. The *asymmetry of the relative phase* is the difference between the relative phases, *ϕr* − *ϕl*. If the eruption of a tooth were simultaneous with both its neighbours, the average relative phase would be 0 (or equivalently 1). If the eruption were a half cycle out of phase with both neighbours, the average relative phase would be 0.5. Because it’s an average of left and right phases, sheering all tooth location columns relative to their neighbours by the same amount does not change the average relative phase. Thus, despite the asymmetry, nearly all dots in panel C are close to 0.5 (dark red). The asymmetry of the asynchrony can be seen in green and purple (Fig. 4C). On the left side of the jaw, left neighbours are shifted slightly forward in time (downward) and right neighbours are shifted back in time (upward) which generates a negative asymmetry score (green). The opposite is true on the right side of the jaw (positive asymmetry score, purple). The centre of the jaw (position 0) has a band of nearly white teeth showing local phase symmetry. The asynchrony and asymmetry observed for Plucker A is evident in several other individuals (Fig. S1, S2).

When we look more carefully at the plucked region in Plucker A, all teeth in locations 15 through 22 that were present in the dental lamina were removed at day 50 (Fig. 5A, box). We were careful not to disturb the free margin of the lamina, thereby ensuring that new teeth would continue to form subsequent to the surgery. This perturbation was designed to test whether already-formed teeth in the lamina might be signaling back to the tooth field at the free margin of the lamina, an idea encapsulated in Osborn’s theory of zones of inhibition. There were wider confidence intervals attributable in the time since last eruption in the surgical site compared to the control side. Although the eruption pattern was disrupted, with some teeth appearing at unexpected times around day 100, by day 150 the pattern was not only re-established but it was in phase with the surrounding unperturbed tooth locations, as reflected in the nearest-neighbour diagonal lines (Fig. 5A). This suggests that the removal of formed teeth from the lamina did not substantially influence the timing of subsequent tooth formation, in contrast with the assumptions of Osborn’s theory. After the surgery the polygons are not as regular (Fig. 5B). The tooth in the centre is out of phase compared to the neighbours (more yellow, Fig. 5B. The position of the centre spot is also offset. The centre tooth erupts much closer in time relative to the immediate neighbours (closer together). The local phase asymmetry illustrates polygons with mirror image antisymmetry (Fig. 5C). Generally, the most sensitive measure of changes due to surgery is the local phase asymmetry. Individual teeth after the surgery are green within the otherwise purple right side of the plot. The colour returns to purple at 150 days or about 90 days since the surgery was carried out (Fig. 5C). These data on local phase asymmetry demonstrate the resilience of the gecko dentition to a major loss in successional teeth.

### Test of the Osborn Model on the plucked animals

We explored possible 3 other mechanisms underlying the observed spatiotemporal patterns in the plucked animals. The first classic Osborn Model, is a literal implementation of Osborn’s theory of zones of inhibition (based on Fig. 3,4 no fixed position for tooth initiation sites). In this model the process of tooth initiation occurs via a diffusing inhibitor, such that the zone of inhibition is spherical, once the tooth expressing the inhibitor has erupted, the dental lamina on either side is no longer inhibited and thus is free to develop teeth. The prediction is that removing teeth, particularly those mineralizing teeth that had migrated out of the tooth field would disturb the pattern since inhibitors would be absent. As teeth erupt and as the jaw grows anterior-posteriorly, space is created for the initiation of additional teeth.

Here we see that there are many inconsistencies between the biological data and Osborn’s Model (Fig. 6). The treated side shows that there would be simultaneous eruption of neighbouring teeth which was not seen in vivo (Fig. 6A). There is no longer a gradient of eruption period (Fig. 6A). The phase relative to neighbours has unrealistic areas in the untreated side where there is no tilt (white circles) and missing central teeth in some of the polygons (Fig. 6B). On the treated side there is no recovery of phase relative to neighbours after the surgery as there is in vivo. The local phase asymmetry is highly abnormal (Fig. 6C). There are large areas spreading out from the midline in which there is no phase asymmetry (white circles). On the treated side there is mixed phase antisymmetry between the right and left tilts which did not persist in vivo.

**Figure 6.**
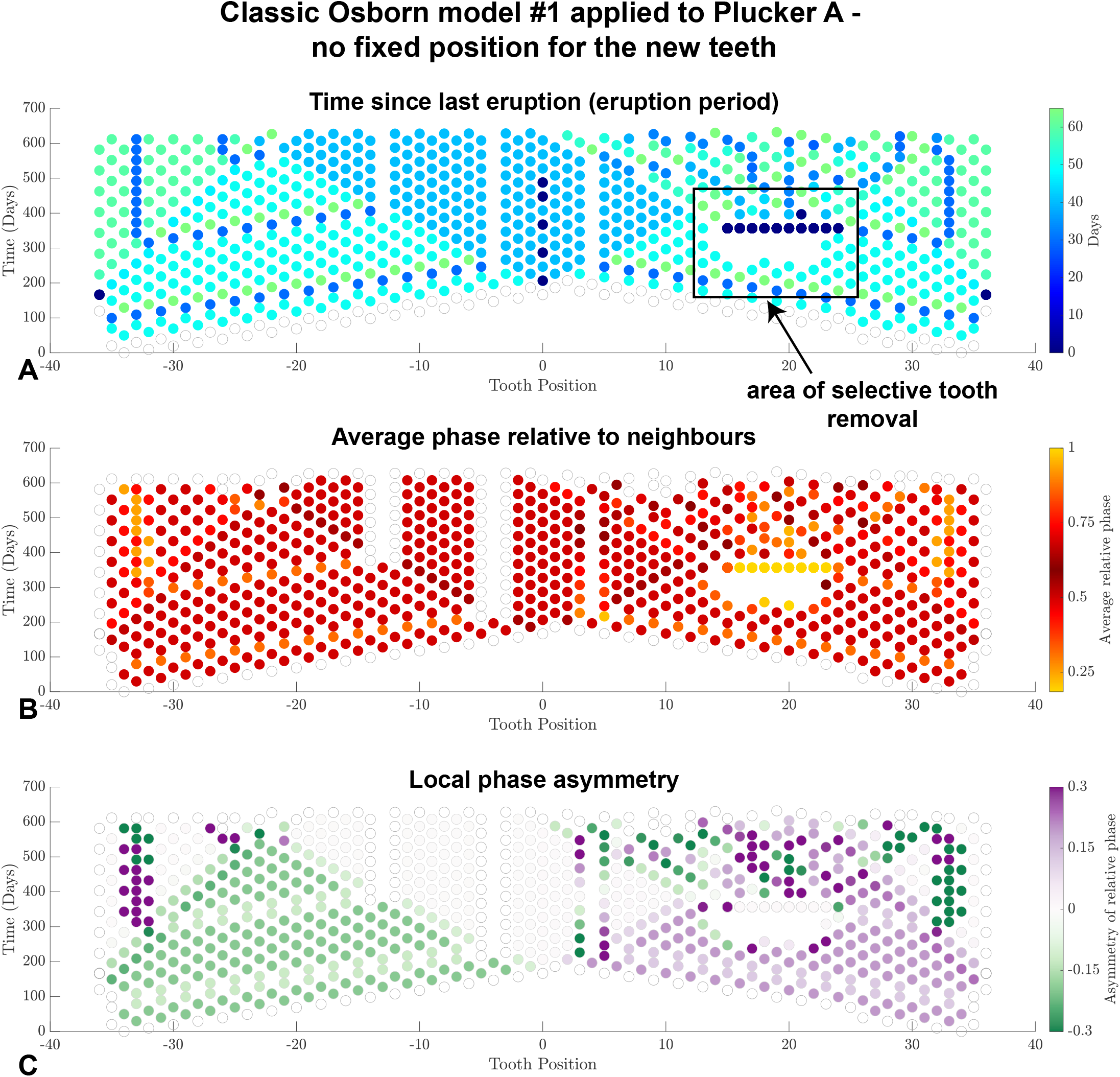
Osborn’s classic model with teeth initiating in positions with lowest level of inhibition. A) The full assumptions of Osborn where perfect circles of inhibition form around each successional tooth was applied to a theoretical plucked animal. New teeth initiate at the intersections of the circles where inhibitor factors are the lowest in concentration. There are many odd patterns of eruption period that form over time including patches with log periods separated by bands where teeth are absent. Most significantly is the area of selective tooth removal. Here the inhibition provided by successional teeth is removed because the teeth have been plucked. When recovery happens, the adjacent successional teeth erupt at the same time which does not happen in vivo. There is no return to presurgical patterns of eruption. B) The average phase relative to neighbours shows the extreme phase shift in the surgical area. There are also bands of teeth that do not appear to have any phase shift (white circles) unlike the in vivo data. C) Local phase asymmetry shows the antisymmetry but there are large periods where there is no asymmetry (white circles). Again this does not resemble biological data. For these reasons the classic model of Osborn is rejected.

We went on to consider a related model in which there are equally spaced predetermined sites along the tooth field at which teeth can form (Fig. 7). In this predetermination model, migrating teeth inhibit any tooth nucleation sites within a fixed radius of their current position. However right from the start, this model also has inconsistencies with the data as shown by a failure of teeth in a tooth family (vertical lines) to form precisely at the lowest level of inhibition (Fig. 7, teeth do not form at the precise intersection of circles of inhibition). The diagonal zahnreihen now connect adjacent tooth positions which does not fit the alternating tooth shedding pattern seen in the real data. In the test of Osborn model #2 there is no evidence of gradual slowing of the rate of tooth eruption (Fig. 8A). There are instead chevrons on very rapid tooth eruption cycles spreading from the midline. These values are not in line with the in vivo rates (Fig. 5A).

**Figure 7.**
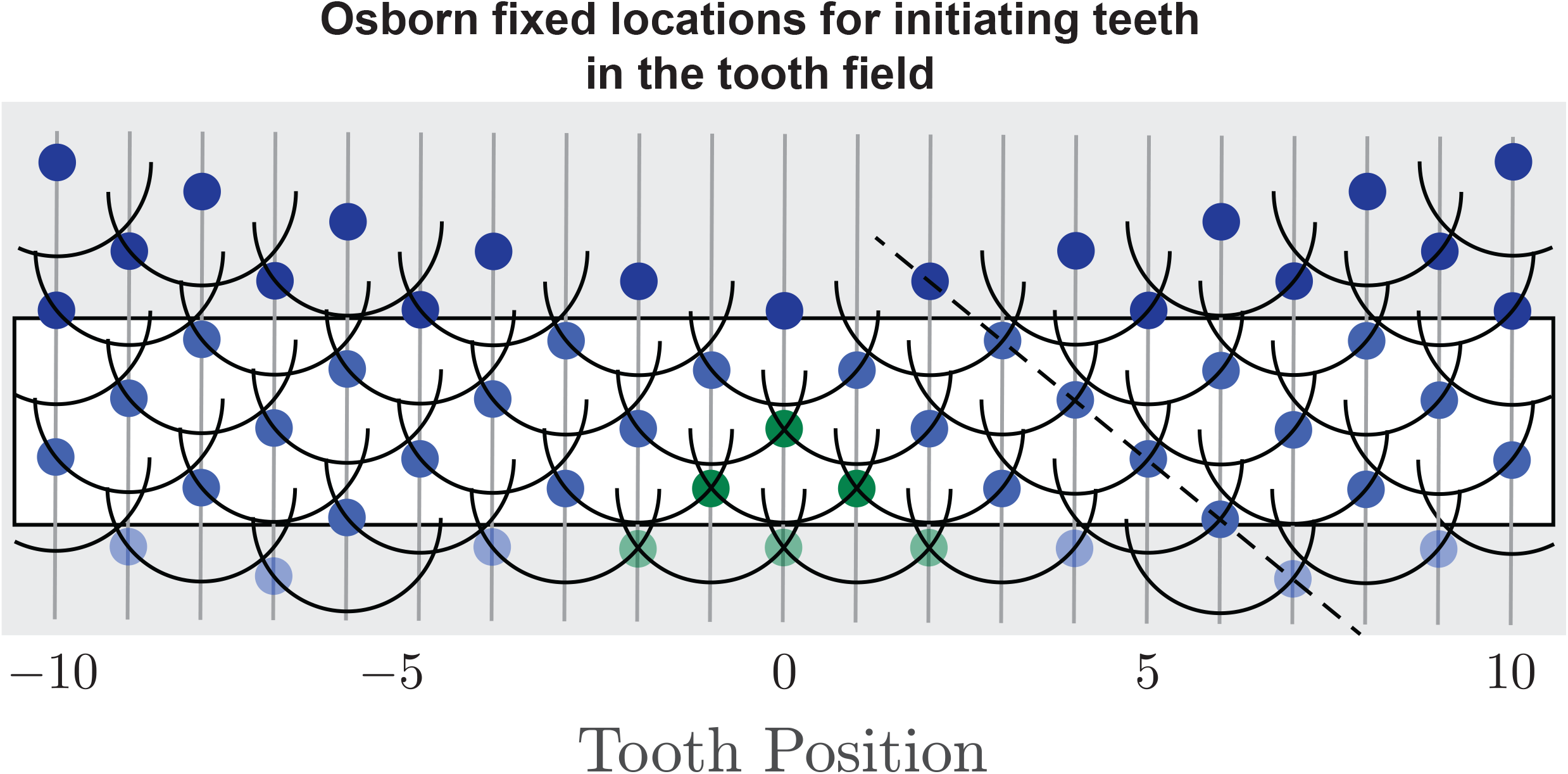
Osborn’s model with fixed locations for tooth initiation. The Osborn model has been adjusted so that teeth initiate in fixed positions in the tooth field. In this representation, the tooth successors are in straight vertical lines without tilt. However now teeth are not initiating at the intersections of circles, possibly outside areas with high levels of inhibition.

**Figure 8.**
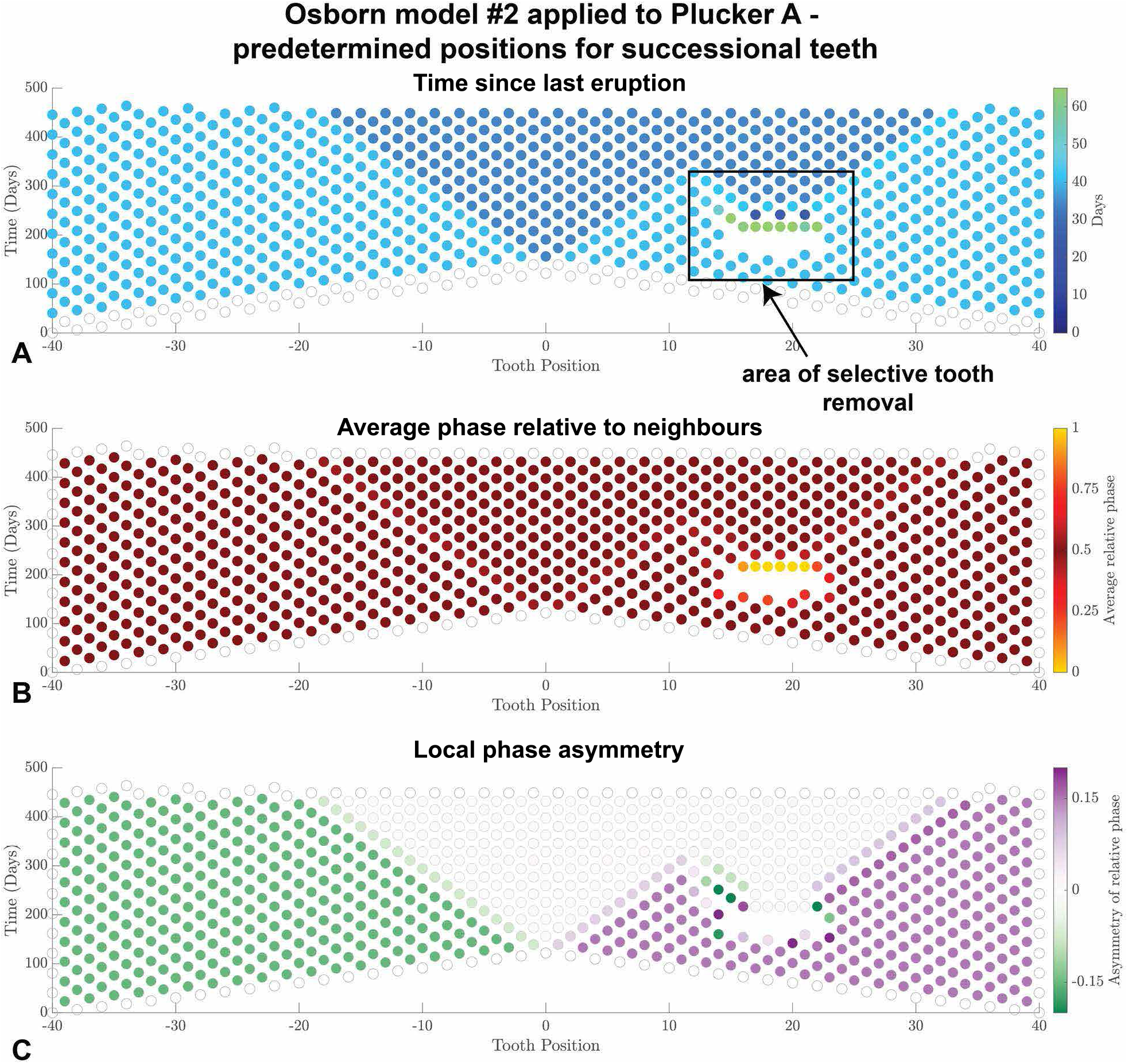
Application of Osborn’s model assuming fixed locations for tooth initiation. **A)** The time since last eruption contains chevrons of consistent eruption times but these do not increase according to increased age of the animal. There are shorter eruption times close to the centre (darker chevron) but gradually more and more teeth are included in this chevron, possibly including the entire dentition. In the area of selective tooth removal there are neighboring teeth erupting at the same time after a delay. B) The relative average phase is very regular on the untreated side with offsets similar to in vivo data. However, the surgical area shows a sharp increase in average phase in the teeth erupting after the surgery. There is no disturbance around the plucked site. C) In the local phase asymmetry analysis is a lack of asymmetry in the centre that extends posteriorly. The treated side shows very minor disruption in phase asymmetry and then loss of the asymmetry which does not happen in vivo. Importantly the tilt is still not sustained and tooth eruption converges to a symmetric pattern. The surgical defects also propagate into a zone where no relative phase asymmetry is present. Therefore, it is unlikely successional teeth initiation in strict vertical positions.

### Variations of Osborn’s model -The Phase Inhibition Model

Next, we considered a third model to more directly capture the biology. This Phase Inhibition Model, assumes that there are fixed nucleation sites, as in the Predetermined Osborn Model but that each site has a natural cycle capable of generating a new tooth every 30 days (Fig. 9). In addition, at some fixed phase in the cycle, groups of cell at that site secrete an inhibitor that slows down, but does not completely inhibit the cycles of neighbouring sites. The partial inhibition causes neighbouring sites to lag by a small amount (Fig. 9). The phase model simulates tooth formation based on a signal emitted from each tooth position. The responding teeth are either sensitive or insensitive to the inhibitor to varying degrees. The time between successive teeth in a single tooth position is based on an ordinary differential equation, which takes into account parameters for the period gradient in time and space along the jaw.

**Figure 9.**
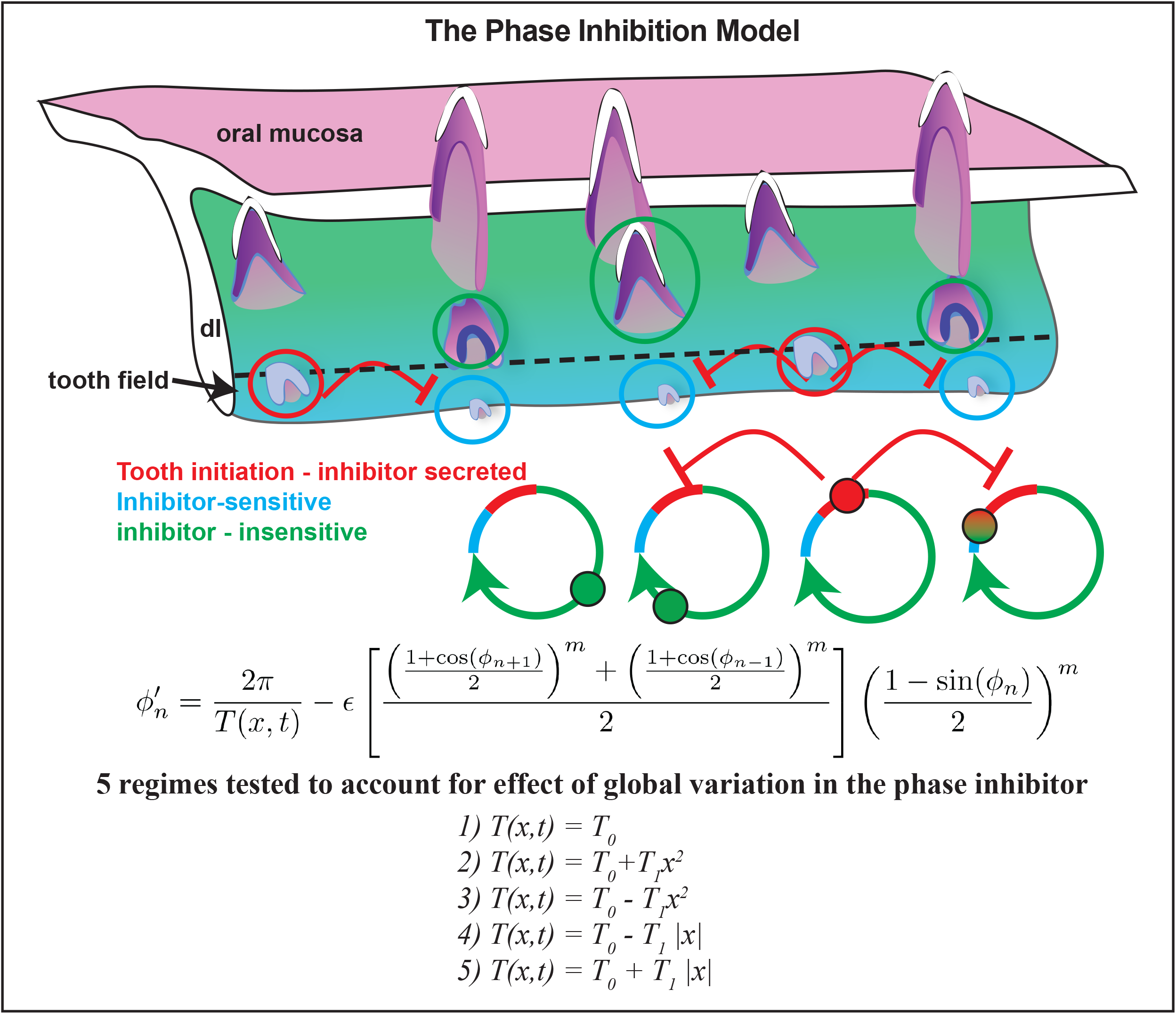
The phase inhibition model. The phase inhibition model assumes there is a predetermined initial configuration for the teeth and there is inhibition from neighboring teeth. Each tooth is entering a cycle of development (circles) and at one particular phase an inhibitor is secreted (red lines, red tooth in the tooth cycles). Teeth cycle through intervals where they are sensitive (blue) or insensitive (green) to the inhibitor. Inhibitors are secreted from bud stage teeth in the tooth field and these are secreted laterally to pause tooth development of the newest tooth buds. The differential equation contains terms to slow the phase growth, thus inhibiting the formation of teeth. The model includes terms that cause inhibition to occur when a tooth is close to forming. This model, unlike the phase model teeth can respond to noise and there is communication between teeth across the jaw.

The differential equation is defined as follows: The cos terms act to slow the phase growth, thus inhibiting the formation of teeth, and the inclusion of sin ensures that this inhibition only occurs when a tooth is close to forming (Fig. 9). This model can respond to noise and contains communication between teeth across the jaw. Several variations in the parameters were tested to see which fit the data the best.

The phase inhibition model fits the biological observations made on plucker A (Fig. 10). The eruption period is very regular and there are almost no differences across the jaw which arises from removing variation between tooth shedding events. There is a delay in eruption period between when the surgery was done and the next teeth erupt (Fig. 10A). The average phse relative to neighbours has little variation, except around the surgery site. Here the phase is only slightly out (0.25) and then returns to normal 0.5 phase (Fig. 10B). The phase asymmetry remarkably captures the in vivo patterns. The anti-symmetry on the right and left sides is perfect, except in the region of the surgery. There are select tooth positions that have lost the phase asymmetry but then these recover after the surgery. There are no effects on the adjacent tooth families. In other words the surgery has local effects only in the phase-inhibition model.

**Figure 10.**
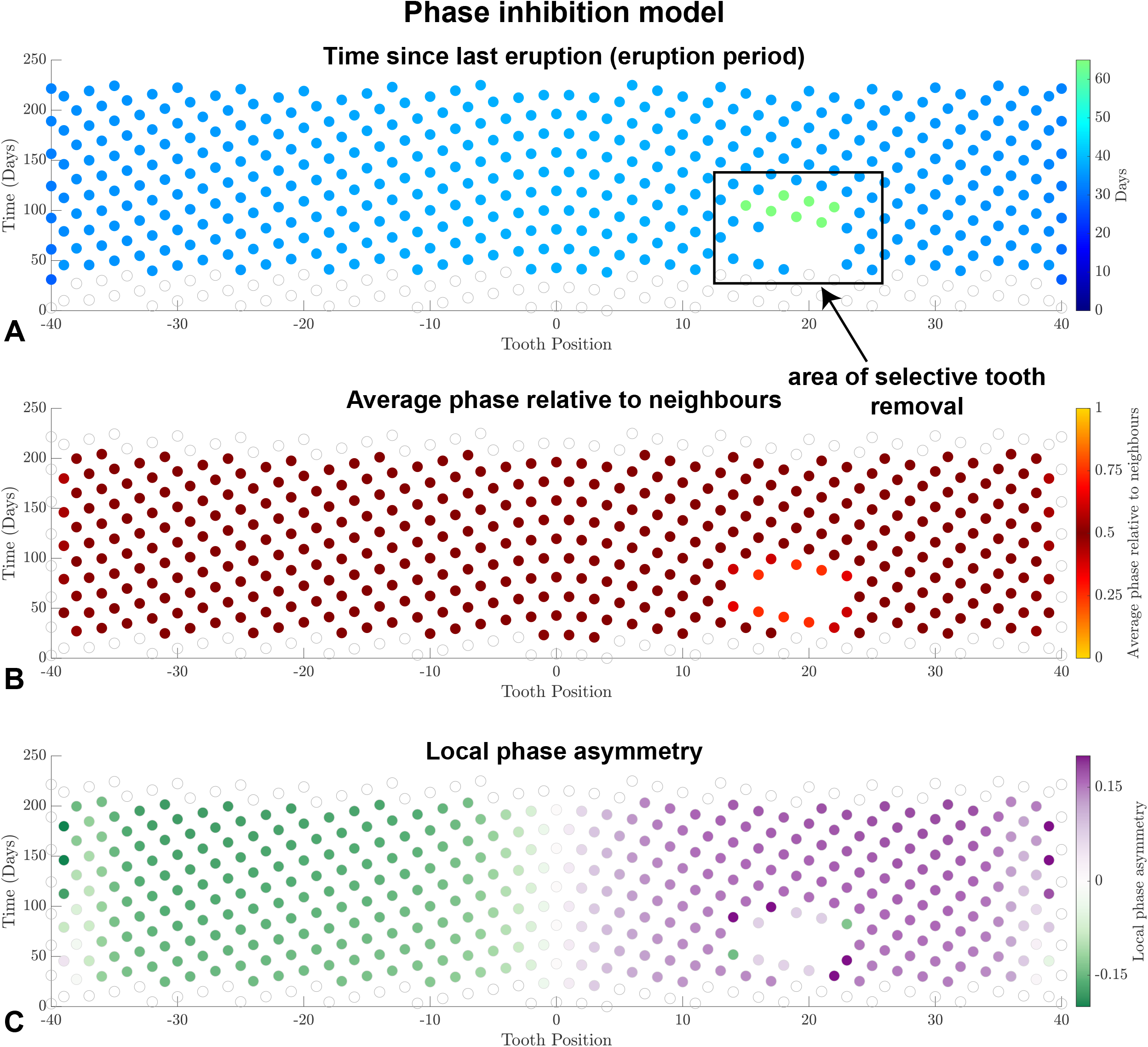
Implementation of the phase inhibition model. A) The model smooths the data, removing variation between tooth shedding events. The same patterns of delay in eruption followed by recover are seen. However the posterior teeth seem to erupt more rapidly than central teeth which is opposite to in vivo data. B) Average phase is also smoothed out and very closely represents the phase distortions caused by plucking. C) The overall mirror image is maintained in the model and the midline is lacking antisymmetry (white circles). The phase inhibition model most closely fits the biological observations made on Plucker A.

### Dental lamina ablation prevents healing after tooth plucking

We carried out experiments that prevented teeth from regenerating in a specific region of the jaw in order to test whether there were local effects at the edges of the region and whether there were any long range effects on tooth replacement. The first set of animals were studied after shorter intervals to confirm the teeth were removed, to examine the effect on dental tissues of applying a solution of 15% Fe_2_(SO_4_)_3._ The animals used for microscopic studies were first scanned with μCT prior to processing (Table 1).

The initial response to application of Fe_2_(SO_4_)_3_ and selective tooth removal was to cause disruption of the dental lamina and an inflammatory response in the mesenchyme. The identification of dental remnants was difficult at this early time point (data not shown) so we concentrated on 2 weeks, 1 month, 3 months and older animals. The 2-week animals had evidence of missing second generation teeth (N= 2; Fig. 11A,B) and a torn dental lamina (Fig. 11B). There were actively proliferating cells in the remnants of the dental lamina (Fig. 11C). There was no PITX2 expression in the dental lamina which is consistent with earlier data showing that after surgery, only the deeper parts of the dental lamina near the tooth field express PITX2 (Brink et al. 2021). We published earlier that the dental lamina heals together between 2 weeks and a month after surgery(Brink et al. 2021). We had therefore expected there would be a dental lamina regenerated. However, after 1 month there was no evidence of teeth forming in the treated area (Fig. 11D). The dental lamina epithelium was extremely short in the treated region (Fig. 11E; N = 2). We checked to see whether proliferation was still taking place and found PCNA staining close to the basement membrane of the dental lamina (Fig. 11F, F’). We also found some cells positive for PITX2 which is ectopic and not seen in the plucked animals (Brink et al. 2021). We also performed a pulse-chase with BrdU to identify whether there were label retaining cells present in the truncated epithelium. Indeed, there were a small number of these cells that took up the label right after surgery similar to dental lamina in animals whose teeth were plucked but not treated with FeSO_4_. At 3 months (n=2) we began to see extensive bone deposition in the area where teeth had once been (Fig. 11H). Interestingly PITX2 expression is present in the basal layer (Fig. 11J’). By 14 months post treatment there was a large amount of bone deposited almost filling in completely the areas where teeth had been (Fig. 11K,L). The dental lamina was either truncated completely (Fig. 11L) or a small tongue of epithelium was present in the treated area (Fig. S4C). Proliferation in the basal layer was maintained (Fig. 11M, N) and PITX2 expression was present (Fig. 11O). We carried out a pulse-chase experiment in Ablator M where the BrdU was administered one month prior to euthanasia but 13 months after surgery. There was uptake of BrdU in the epithelial cells (Fig. 11P) so putative stem cells could still be present even in the very short dental lamina. Despite having these label-retaining cells, no extension into the deeper tooth forming regions occurred. Interestingly, iron particles were retained in the mesenchyme (Fig. 11E,I,L).

**Figure 11.**
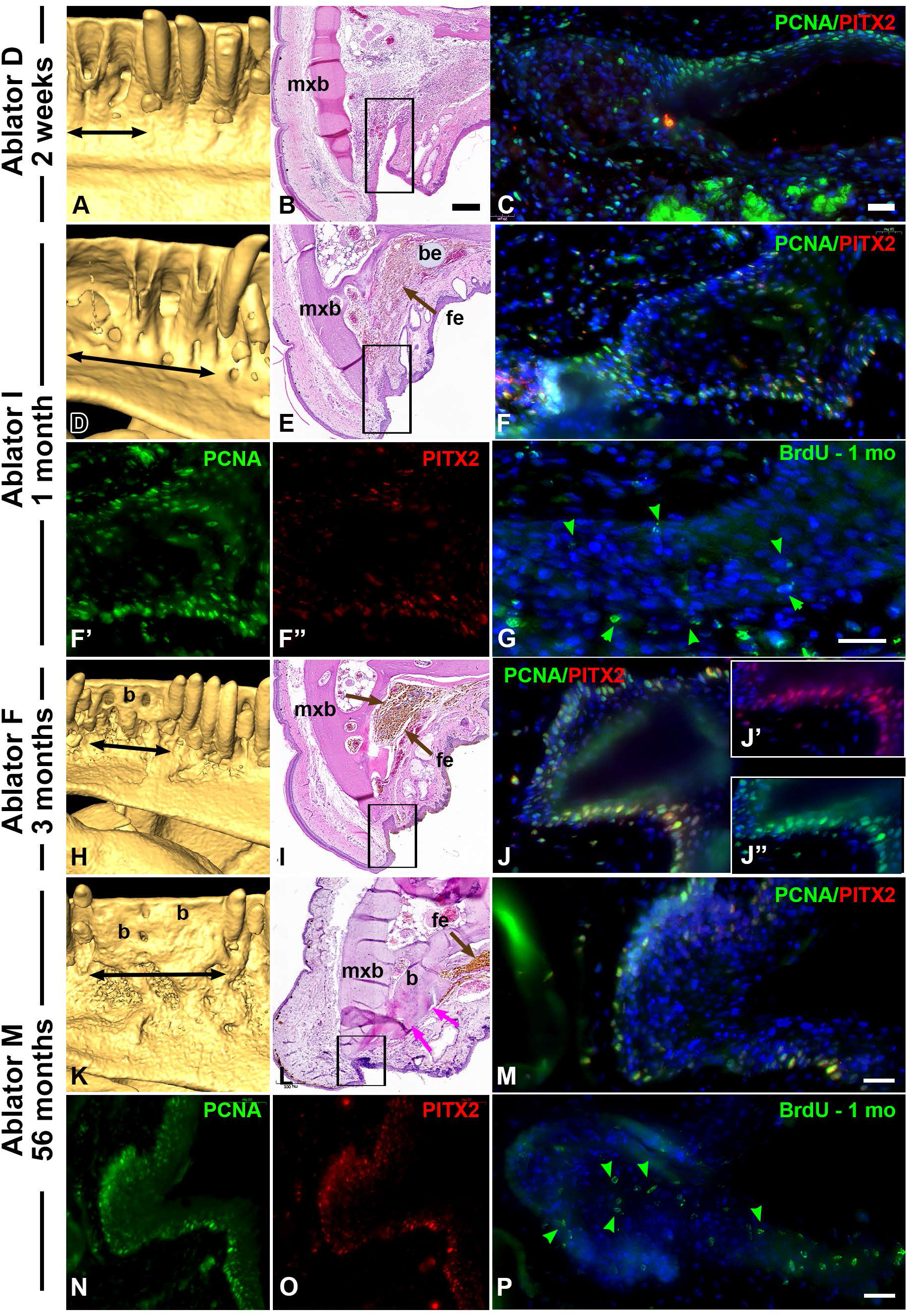
Effects of local chemical cautery with Ferric sulfate on the replacing dentition. The full pipeline of analysis includes μCT after fixation of the jaws, decalcification, embedding in paraffin and histological analysis. A-C) Two weeks after ablation, a torn dental lamina is visible and proliferation is present in the basal layers of the dental epithelium as shown with PCNA staining. There is no PITX2 staining. D-G) one month after surgery, bone has started to fill in and the functional teeth have not returned in the ablated area (double ended arrow). E) In histology a short dental lamina is visible (box) and iron particles are in the mesenchyme where the tooth field used to be located (brown arrow). An affigel blue bead was implanted to mark the surgically treated area. F-F’) The short dental lamina has basal epithelial cells that are proliferating as shown by PCNA staining. There are some cells that are PITX2 positive. G) an adjacent section from the same block showing retention of BrdU label, one month after the initial pulse. Most the cells are likely quiescent label-retaining cells (green arrowheads). H-J’’) At three months more bone has filled in the gap which prevents new teeth from growing in this region. I) a very truncated piece of dental lamina is present near the oral surface (box) as are iron deposits in the mesenchyme. J-J’’) The proliferation of the truncated dental lamina is confirmed and there is PITX2 staining in the same basal cells (J’). K-P) This animal was followed for 56 months. The bone has completely filled in the places where functional teeth used to be present (K). in sections a short dental lamina persists (L). Basal cells are proliferating (M, N) and some of the cells express PITX2 (O). The pulse chase experiment carried out one month prior to euthanasia labeled some of the epithelial cells (green arrowheads). Label retaining cells appear not to overlap with PITX 2 or PCNA stained regions. Key: b – bone apposition, be – bead, fe – iron, mxb – maxillary bone. Scale bar in B = 200 μm and applies to E, I, L. Scale bar in C = 20 μm and applies to F-F’’, N,O. Scale bar in G = 50 μm and applies to J-J’’. Scale bar in M = 20μm and applies to M, P.

We wondered whether the iron particles themselves may have inhibited tooth formation so we examined adjacent teeth that were not part of the selective tooth removal (Fig. S3A,A’, B, B’, C, C’, D, D’). There were iron deposits extending into adjacent regions (Fig. S3A’,D’). However, there was normal proliferation in the cap-stage teeth next to the iron deposits as shown with PCNA antibody staining (Fig. S3A’’-D’’). Moreover, the pulse of BrdU had been retained by some epithelial cells at 1 month (Fig. S3B’’’.D’’’) and 3 months after the initial pulse (Fig. S3C’’’). The iron deposits themselves are not a barrier to forming teeth. We presume that it is the absent dental epithelium within the ablated zone that is likely the cause selective tooth agenesis.

### Long term tracking of the dentition after ablation

We next examined 3 animals that had single locations of ablation in one quadrant of the maxilla and followed them for up to 9 months with wax bites. We hypothesized that if there were diffusible inhibitors or activators then effects on adjacent tooth families outside of the ablated region would be seen. Moreover, if there was unidirectional signaling then only teeth posterior or anterior to the treated site would be affected.

### Testing the Wave Stimulus model of Edmund on the ablated animals

In Edmund’s wave stimulus theory (Edmund 1960; 1962), an extrinsic chemical signal originates from the first tooth position and disperses distally along the jaw, inducing the initiation of tooth development as it proceeds. As a result, stimuli propagate along the jaw and are responsible for subsequent generations of teeth. The forward-temporal staggering of tooth eruptions initiates a lattice dentition pattern. This theory relies on two key assumptions: the centre tooth is the first to develop; and teeth are progressively initiated in order along the length of the tooth row. There may be predetermined sites of tooth formation. The ablation of the tooth field is predicted to make the signals weaker or abolish them entirely. Alternatively, The signals may be propagated from tooth family to tooth family. Ablation of the tooth forming field should weaken these signals, especially close to the surgery site.

The data from Ablator M (Fig. 12 and S4) shows a gap in tooth replacement that resolved to an absence of 6 tooth positions. Ablator L shows a similar gap in about 6 tooth positions on the treated side which was confirmed in contrast-enhanced μCT (Fig. S4A-C). In ablator M the time since eruption was stable on the untreated and treated sides of the jaw (Fig. 12A). There were some teeth that erupted more rapidly than normal adjacent to the surgical site. The average phase relative to neighbouring teeth was minimally disturbed with most differences being located bordering the posterior edge of the treatment site (Fig. 12B). The analysis most sensitive to directly changes in tooth replacement was local phase asymmetry. However, the right and left sides were still largely anti-symmetric or mirror image (Fig. 12C). There were only a few teeth that appeared to be asymmetric and these were immediately posterior to the surgical site.

**Figure 12.**
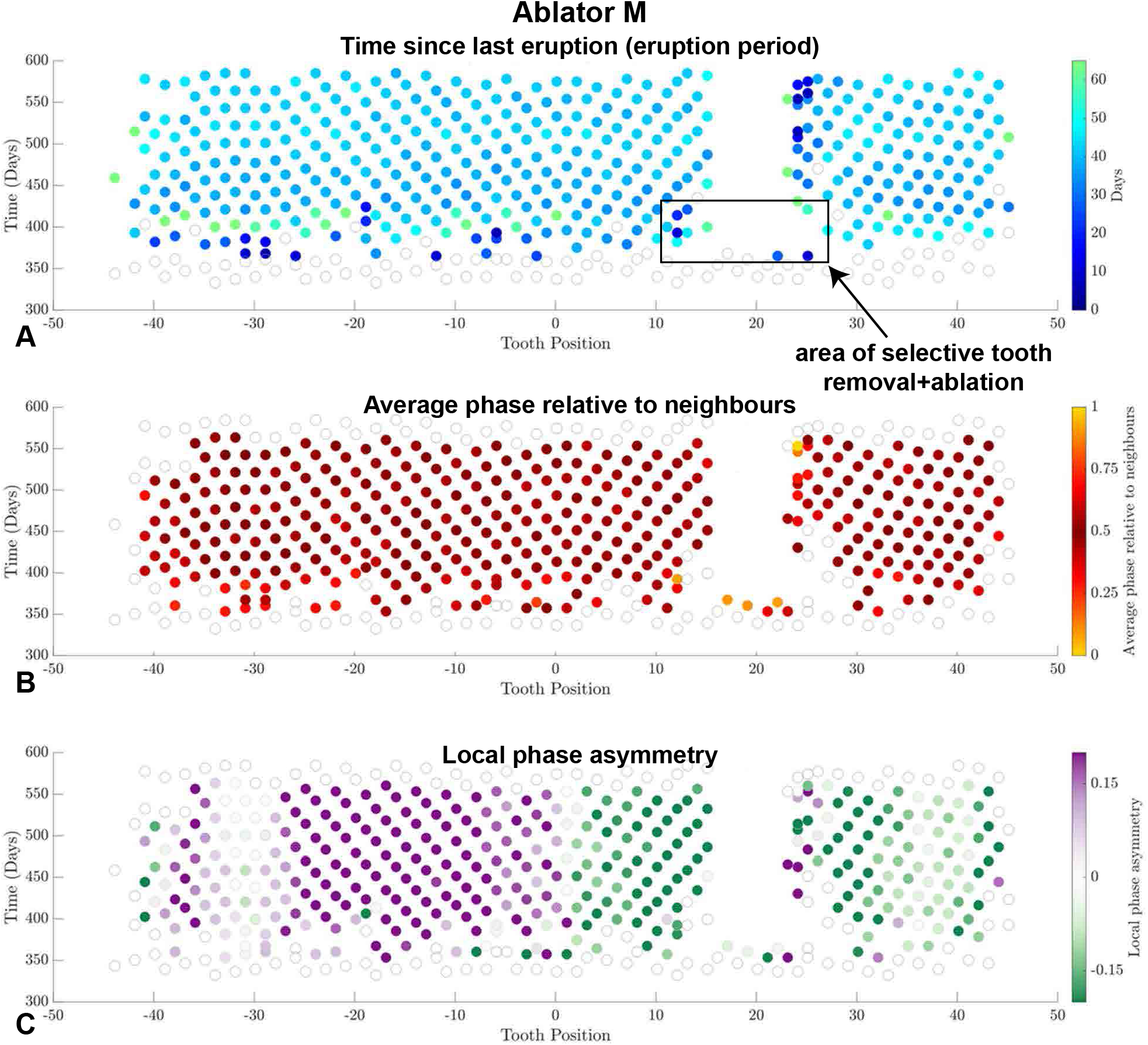
Effects of ablation on long term tooth replacement patterns. A) The application of Ferric Sulfate resulted in the permanent loss of approximately 6 tooth positions. There is minimal disruption of eruption period except for teeth directly adjacent to the treated site. B) The phase relative to neighbours remains at 0.5 in most tooth families. Only those bordering the area posterior to the surgical site have acquired a phase that is closer to 1 or zero. C) Local phase asymmetry analysis shows minimal disruption across the surgical site supporting the idea that there is no long range signaling taking place across the gap.

Edmund’s wave stimulus model does not work with our data in ablated geckos. Instead of a totally independent signal that propagates across the jaw or even one that is transmitted from one tooth family to the next, our data shows only local signaling within tooth families is needed to maintain jaw-wide patterning. Furthermore, there is no evidence that the signal weakens after tooth ablation.

## Discussion

We have uncovered striking patterns of tooth replacement in leopard geckos that can be used to derive models of tooth replacement. We know that the period of eruption slows in time and along the jaw. Tooth shedding is remarkably anti-synchronous. There is a slight tilt or off-set that is opposite on the left/right sides (green/purple) of the jaws. We have captured nearest and second nearest neighbours patterns which were proposed many decades ago in comparative studies (Edmund 1958; Edmund 1960; Woerdeman 1919). Finally, in the leopard gecko we uncovered tremendous resiliency. When teeth were plucked out the pattern recovers perfectly after 3 tooth cycles or 3 months.

### Comparison of Osborn’s zone of inhibition relative to the phase-inhibition model

In Osborn’s model there is signaling from teeth that have already developed (second generation teeth) and then when these teeth move towards the oral cavity, the teeth escape inhibition and can complete development. In μCT scans of pluckers, we never saw teeth of an advanced stage deep in the tooth forming field, waiting for a space in the inhibitory zone to appear (Brink et al. 2021). When teeth were plucked, the immature teeth form deep in the tooth field but once they have formed a substantial crown complete with the enamel cap, these teeth have migrated out of the tooth field. There are never teeth of the same stage beside each other, but neighbouring teeth are arranged in diagonal rows or zahnreihen according to maturity(Brink et al. 2021). We also never saw a group of teeth erupt at exactly the same time which would have happened if there was a simultaneous release from inhibitory signals. Based on our results, we have ruled out inhibitory signals being produced by mineralizing second generation teeth. There is no evidence of downward signaling from oral to aboral tooth field. Instead effects of plucking teeth are locally constrained to neighbouring teeth. Different mechanisms to regulate the patterns of tooth replacement may be involved.

In our Phase-inhibition model, teeth can continue to develop and move towards eruption while not being completely inhibited. The phase-inhibition model also includes periods where the teeth are relatively sensitive or insensitive to inhibitory signals. In the model, there may be one source of inhibitory molecule in the tooth field but teeth themselves alter their cell surface receptor expression to be less or more responsive to the signals. Of all the signals investigated so far, the canonical WNT signaling pathway seems the most promising candidate. The alligator (Wu et al. 2013) and iguana (Brink et al. 2020) successional laminae are enriched for β-catenin protein relative to surrounding tissues. This suggests that canonical WNT signaling promotes tooth initiation. In the alligator there is one WNT inhibitor expressed, SFRP1 protein expressed in the pre-initiation stage and then decreased once teeth initiate (Wu et al. 2013). This suggests that when the inhibitor is downregulated, teeth can begin development suggested by Osborn. When WNT3A was added to cultures of alligator tooth buds the level of proliferation was increased, however SFRP1 protein inhibited proliferation.

More work needs to be done to localize the inhibitors, activators and WNT receptors in the adult leopard gecko. Previously we found that *DKK3* (a stem cell marker and canonical WNT antagonist) is expressed in a small group of cells in the dental lamina close to the budding teeth (Handrigan et al. 2010). In addition, the juvenile gecko has high levels of expression of canonical WNT target genes, *TCF4* and *LEF1* close to the newest budding teeth suggesting the pathway is active at the initiating stage of development (Handrigan et al. 2010). We completed functional studies in organ cultures of gecko tooth buds to test whether canonical WNTs stimulate tooth proliferation. Indeed, adding BIO to the cultures, a drug that activates canonical WNT signaling, increases gecko dental epithelial proliferation (Handrigan et al. 2010). However, in vitro cultures carried out in the gecko and alligator (Brink et al. 2021) do not support tooth succession or 3D morphogenesis so we don’t know whether activating this pathway would alter tooth succession in vivo. In the future we hope to develop a molecular approach to studying tooth replacement that can be used in the highly accessible gecko dentition.

We acknowledge there are some limitations to our study. Foremost is the variability between animals. We have provided the raw data for all three pluckers and even though there is variability, it is still possible to see similar trends such as slowing down of tooth eruption rate over time, conserved average phase of 0.5 between neighbouring teeth and largely anti-symmetrical phase asymmetry in the right and left sides of the arch. There are other reasons for variability in the data mainly related to manual scoring of the data. It is challenging to keep track of missing tooth positions especially right after plucking and in the long term in the ablators where tooth positions are permanently lost. We propose in the future to develop an image analysis application to better account for tooth positions after treatment. Some of the planned steps include finding the midline, straightening the arch perpendicular to the midline and then using machine learning to score the indents. We discovered that the input of raw eruption data into mathematical plots reveals sites with potential errors in scoring. We were able to check the wax bite raw data against the real wax bite data for accuracy and correct the errors..

We have been able to rule out the Osborn model of inhibitory factors regulating the timing of tooth initiation. We have shown in the real data that successive teeth do not migrate laterally over time. It also seems likely that the tooth forming field provides the signals for tooth initiation. We also ruled out two versions of the wave-stimulus model of Edmund using the ablator animals. We can now develop new models of tooth replacement and simulate their effect on the dentition by adding on tooth cycles. These simulations will be used to test and refine future models. Finally, it is normally hard to measure biological resiliency especially with full organ regeneration. Wound healing in the skin does not necessarily restore all the ectodermal specializations and scar tissue forms in the dermis. Gecko teeth offer a unique opportunity to study a complex repeating unit from the point of initiation to full organ function. We are optimistic that that the minimally invasive experimental methods in adult gecko dentitions, combined with sensitive mathematical tools will shed new light on maintenance of patterned organs and tissues.

## Supporting information

Brink et al. Supplementary figures

## Funding

The experimental work was funded by NSERC grant RGPIN-2016-05477 and NIH grant 5R21DE026839-02 to JMR. Funding for the gecko surgical suite was from NSERC grant RTI-2016-00117 to JMR. KSB was funded by a Killam Postdoctoral Fellowship, a Michael Smith Foundation for Health Research Postdoctoral Trainee Award, and an NSERC Banting Postdoctoral Fellowship. TMG was funded by an NIH F32 Individual Postdoctoral Fellowship F32DE024948.

## Conflict of Interest statement

Author Theresa Grieco is employed by the company STEMCELL Technologies. All lab work was performed prior to employment and STEMCELL Technologies had no financial or experimental contribution to this work. The remaining authors declare that the research was conducted in the absence of any commercial or financial relationships that could be construed as a potential conflict of interest.

## Acknowledgements

For assistance with animal care, we thank the veterinarians and staff at the Centre for Comparative Medicine, UBC. We also acknowledge the Centre for High-Throughput Phenogenomics at UBC for conventional μCT scans of the geckos and Tomas Zikmund at CEITEC, Brno, Czech Republic for the contrast-enhanced PTA scan of the gecko jaw. The authors would like to thank those who participated in the wax bite process over the years since this project initiated: Andrew Wong, John Whitlock and Rodolfo Martin del Campo. We acknowledge the input into the wax bite analysis of Lauren Holtzman and Alex Fraser from the Department of Mathematics. Katherine Fu assisted with some of the immunostaining.

## Supplementary Figure legends

**Figure S1**. Plucker B has some variation in the pattern compared to Plucker A. A) there is a row of teeth at around the time the surgery takes place that have lengthened periods of eruption on the untreated side. There is a gap present on the treated side and this fills in a disorganized way. It is likely that a longer period of follow up would have been needed to detect full healing. B) the average phase of neighbouring teeth is close to 0.5 on the untreated control side. The treated side shows deviations of average phase within and adjacent to the plucked region. C) Local phase asymmetry shows overall anti-symmetry of the data with the green tooth eruptions being on the left side and purple on the right. There are bands of symmetrical eruption in the midline confirming we correctly identified the midline tooth. After surgery there are periods where teeth are out of phase compared to neighbours. Again this analysis shows that a longer follow up was needed to see recovery.

**Figure S2**. Plucker C was followed for a period of about 250 days. The general patterns of eruption period show lengthening as the animal ages. The disruption after plucking lasts about 3 tooth cycles and then mostly returns to a similar pattern compared to the control side. There are exchanges of teeth that are more rapid than normal post surgery. B) The average phase shows that in some regions after the surgery some teeth are out of phase with their neighbours. There is mostly normal phase relative to neighbours surrounding the surgical site. C) The midline as shown by reflected asymmetry is off by one tooth position. The midline should be at -1. There is a reversal of phase asymmetry on the control side of the jaw indicating this may occur as part of natural variation. On the treated side most of the teeth observe the anti-symmetry purple pattern.

**Figure S3. Section of tissues at the edge of treated areas where teeth in early stages of development**. Brightfield views show iron deposits in the tissues adjacent to the ablation region. Fluorescent images show the proliferation patterns with PCNA antibodies (A’’, B’’, C’’, D’’). The retention of BrdU is shown in a split channel after an initial 1-week pulse is shown in the same tooth buds (green arrows, A’’’, B’’’, C’’’, D’’’). **A –B’)** Two regions from the same animal, ablator F. There are iron deposits next to the budding teeth (A’, B’). PCNA staining shows typical high levels of proliferation in the enamel organ and dental papillae (A’’, B’’). There was no retention of BrdU in a second generation tooth (A’’’). The adjacent tooth family did retain BrdU in the successional tooth bud (3, B’’’). **C-C’’’)** In ablator I, followed for 3 months after surgery, there is a second generation tooth bud with strong PCNA labeling in the tooth bud (B’’) and retention of BrdU in the dental lamina (B’’’). The iron deposits did not affect proliferation. **D-D’’’)** Ablator M was followed for 56 months post-surgery and 1 month after the pulse chase labeling. Proliferation is still occurring as shown by PCNA and BrdU co-labeling (D’’, D’’’’). Key: 1-1^st^ generation tooth 2-2^nd^ generation tooth, 3 – 3^rd^ generation tooth, dl – dental lamina, Fe – iron deposits, mxb –maxillary bone. Scale bars = 200 μm for A,B,C,D; 50 μm for A’,B’,C’,D’; 20 μm for A’’,A’’’,B’’,B’’’,C’’,C’’’,D’’,D’’’.

**Figure S4. The raw scoring data for Ablator M**. All wax bites were scored from photographs. The midline was identified and positions identified lateral to the midline (red column M). The number of teeth originally removed and treated with Ferric Sulfate was from positions 12-23 in the centre of the left maxilla. Surgery took place on March 29, 2017 and wax bites were completed on October 30, 2017. The eventual tooth positions affected was fewer than treated, 15-21. This is due to the spread of liquid Ferric Sulfate. The black boxes indicate teeth that were missing, white boxes are teeth present and grey boxes are partially erupted teeth. Although patterns are visible in tooth eruptions it is difficult to measure the phase relative to neighbouring teeth in the raw data. B,C) The same animal scanned after euthanasia on May 29, 2018 showing how bone has filled in the tooth forming region on the left maxilla. The replacement teeth are forming as normal anterior and posterior to this region.

**Figure S5**. Ablator L was treated on March 30, 2017 with plucking and Ferric Sulfate treatment. The animal was euthanized on May 25, 2018. After euthanasia, the jaws were stained with phosphotungstic acid to enhance contrast and then scanned using high resolution μCT (17 μm). A) the volume rendering shows that the surgical site does not have erupted teeth. B) a slice view adjacent to the surgical region shows normal functional teeth, with a dental lamina and a successional tooth. C) In the surgical site there is increased thickness of bone on the palatal side of the maxillary bone. The dental lamina is present but extremely short. D) the wax bite data shows a period where bites were not taken at about 110 days. The right side has a gap where teeth have not erupted and this gap has narrowed from the edges starting at about 225 days. There are likely some errors in scoring the missing teeth which has led to the placement of teeth into the surgical zone (green dots). The overall patterns of eruption can be seen on the control side after 200 days. E) Phase averaging reveals that there are some areas on the control side that may not have been scored accurately. F) The local phase asymmetry shows that the midline is correctly identified as position 0. The same discrepancies in the period between 150 and 225 days are seen on the control side of the jaw.

## Notes

### Competing Interest Statement

TG is currently employed by the company STEMCELL
Technologies. All lab work performed by TG was prior to employment at STEMCELL Technologies. STEMCELL Technologies has no financial or experimental contribution to this work.

