## Supplementary material for "Evolutionarily conserved waves of tooth replacement in the gecko are dependent on local signaling": Brink et al. Supplementary figures

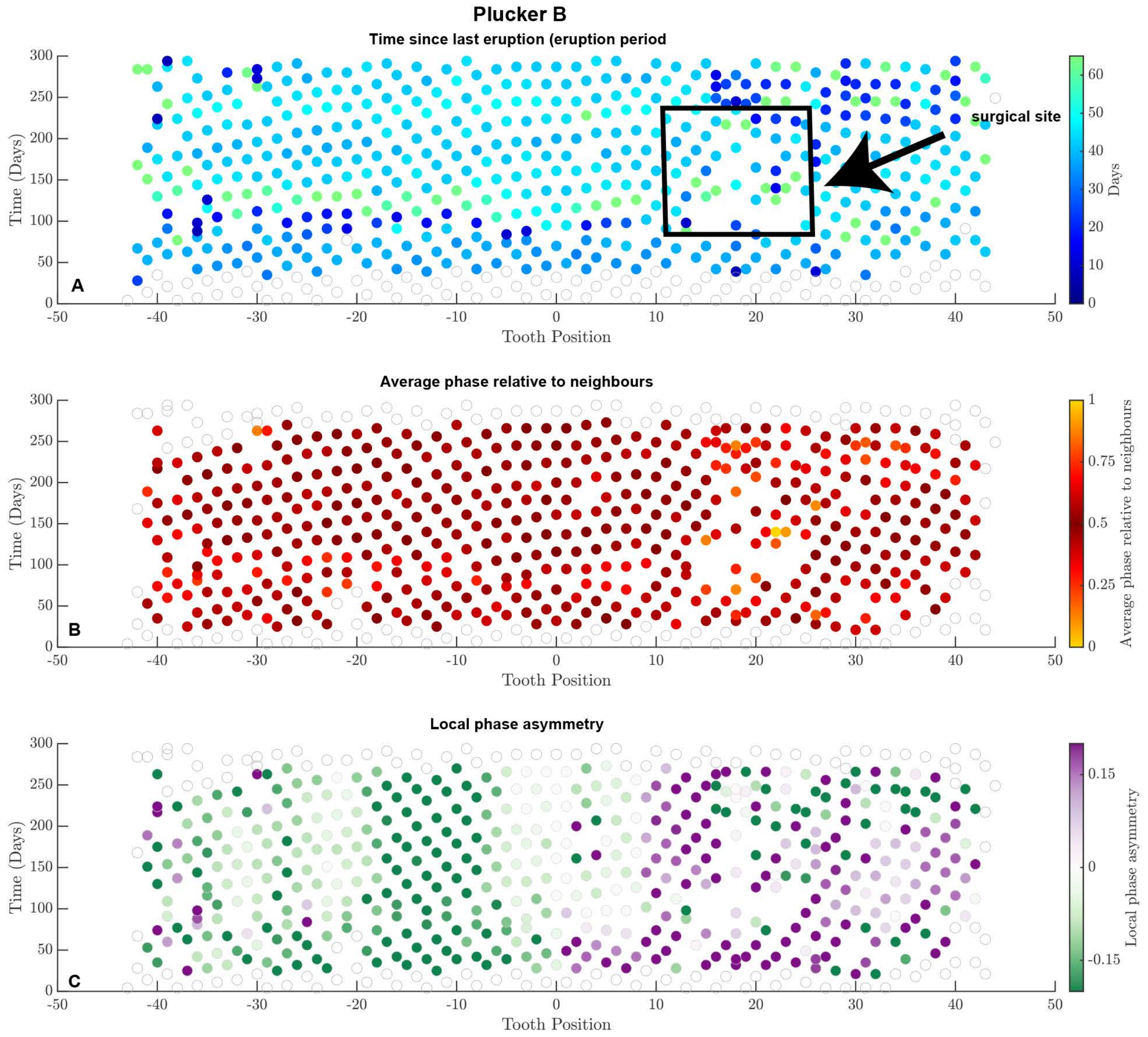
Plucker B
Time since last eruption (eruption period
surgical site
Days
Average phase relative to neighbours
Average phase relative to neighbours
Local phase asymmetry
Local phase asymmetry
Time (Days)
Tooth Position
A
B
C

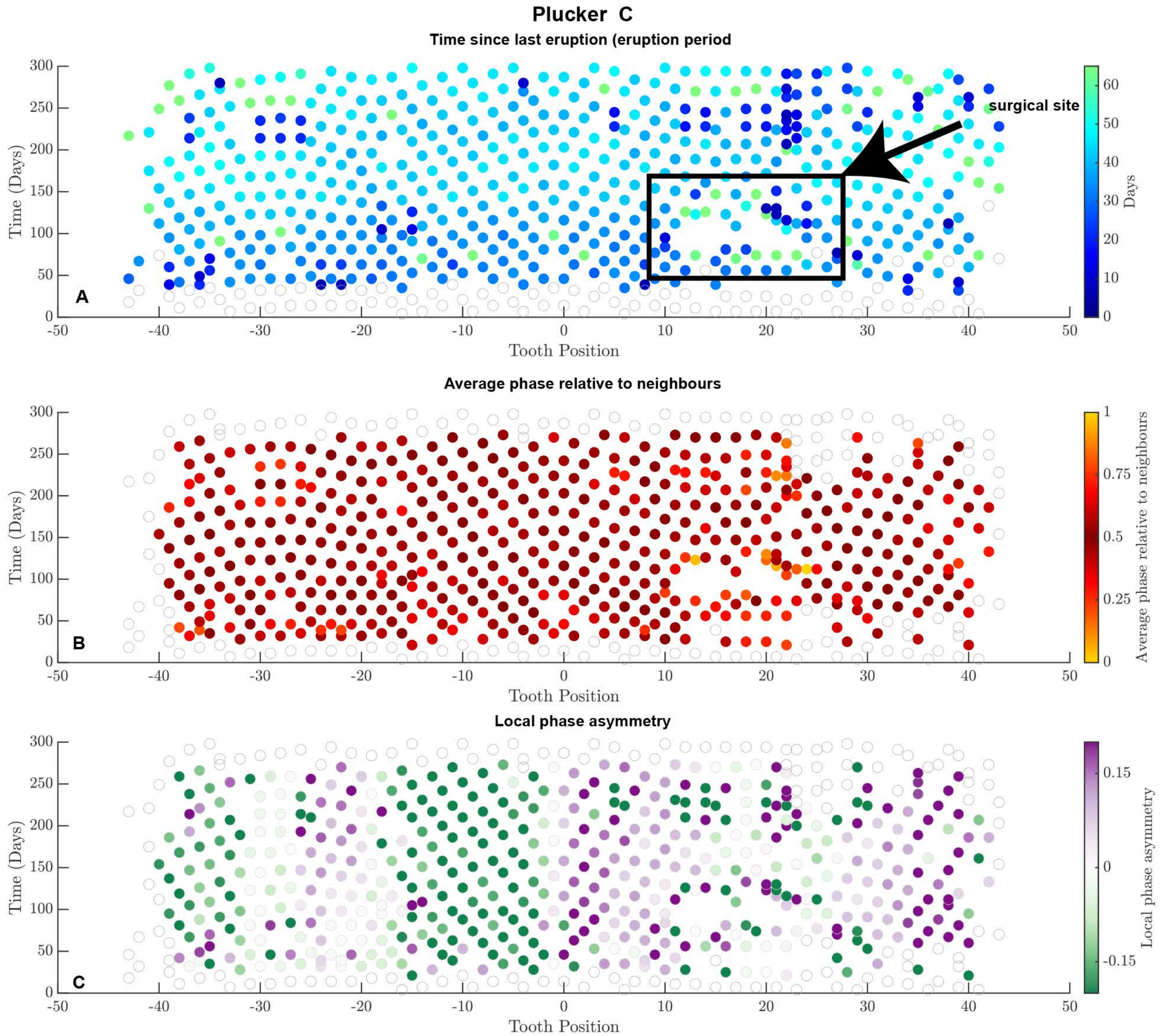
Plucker C
Time since last eruption (eruption period
surgical site
A
Time (Days)
Tooth Position
Days
Average phase relative to neighbours
B
Average phase relative to neighbours
Local phase asymmetry
C
Local phase asymmetry

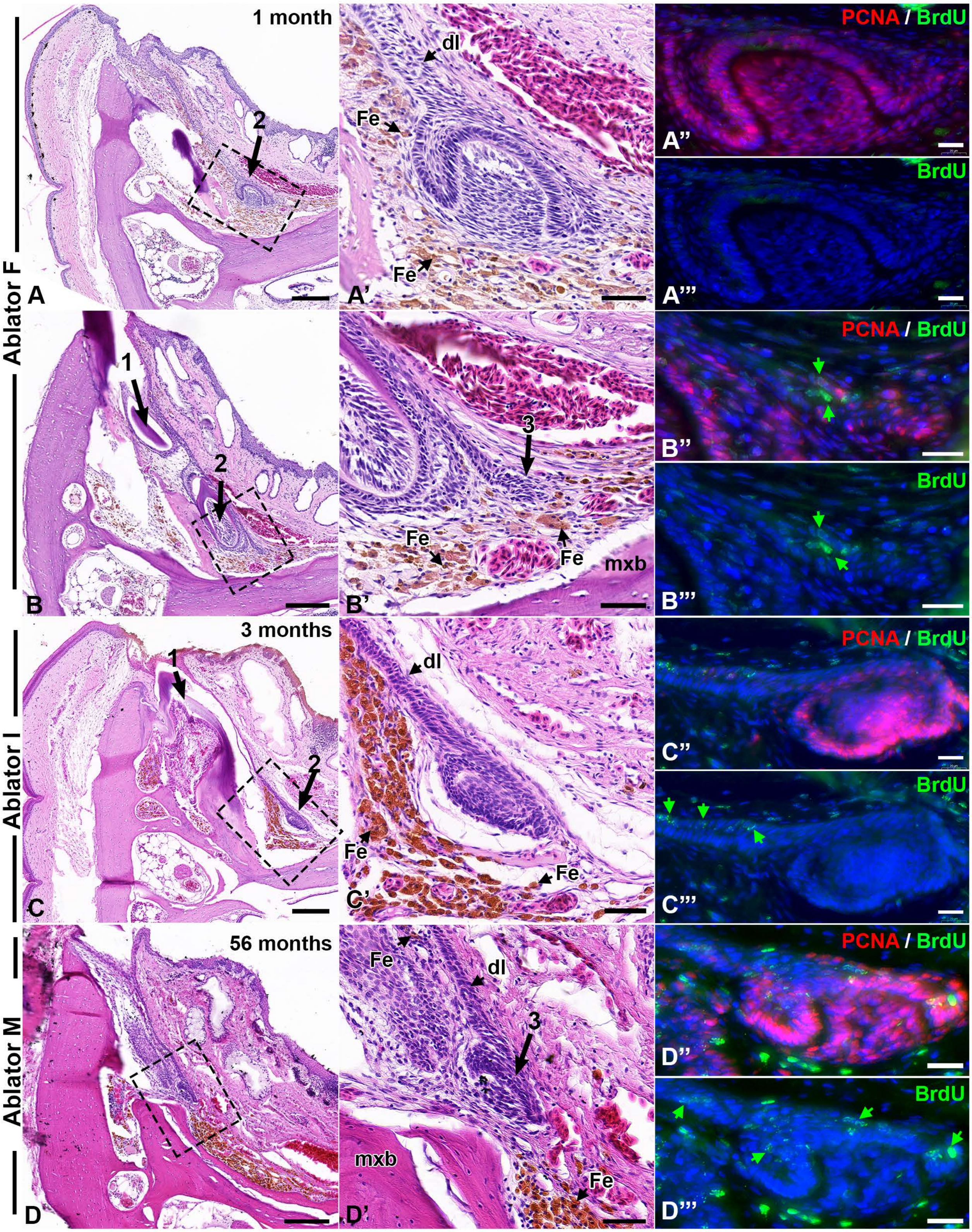

Ablator F
Ablator I
Ablator M
1 month
3 months
56 months
A
A'
A''
A'''
B
B'
B''
B'''
C
C'
C''
C'''
D
D'
D''
D'''
PCNA / BrdU
BrdU
dl
Fe
mxb
1
2
3

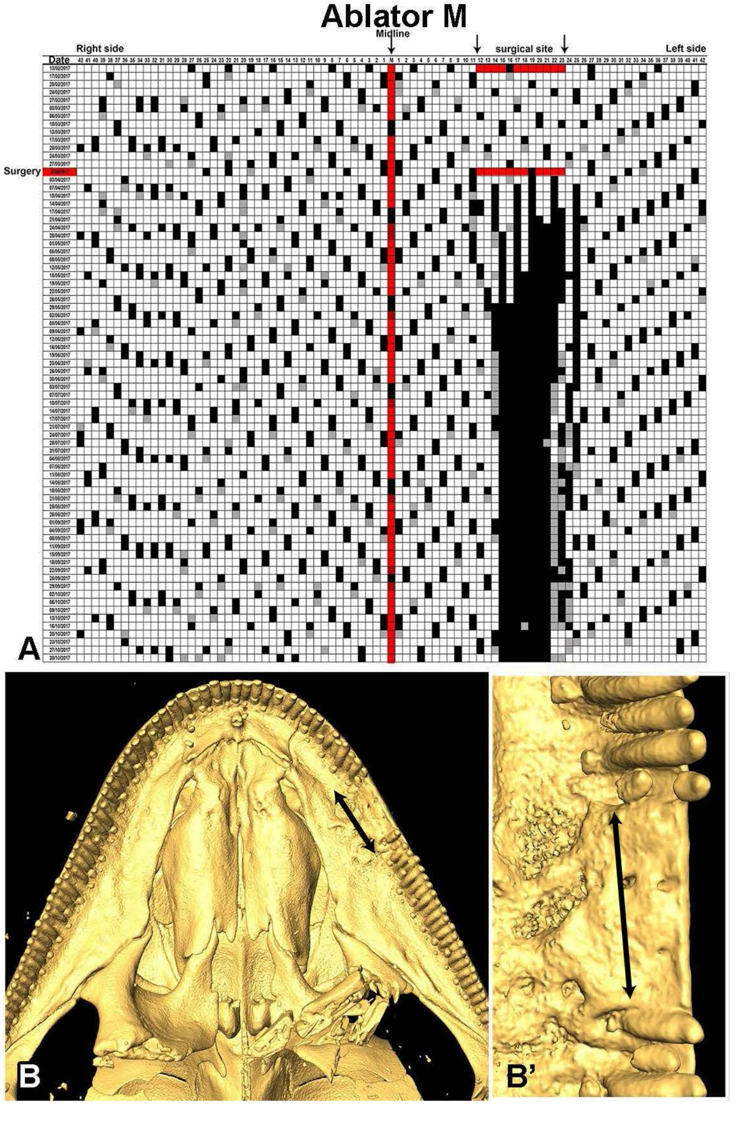

Ablator M
Midline
surgical site
Right side
Left side
Date
Surgery
A
B
B'

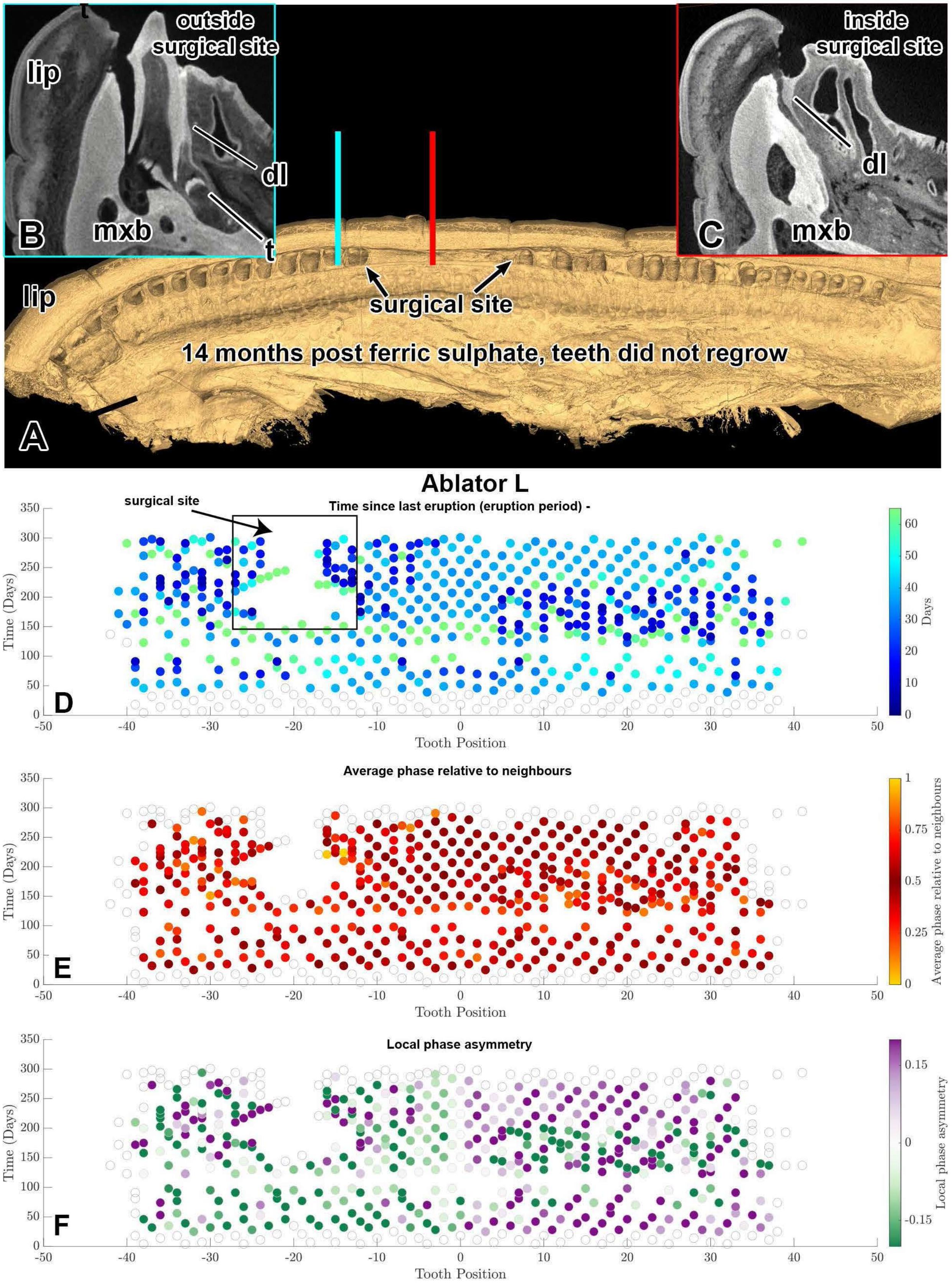
outside surgical site
lip
dl
t
B
mxb
inside surgical site
dl
C
mxb
lip
surgical site
14 months post ferric sulphate, teeth did not regrow
A
Ablator L
surgical site
Time since last eruption (eruption period) -
Time (Days)
Tooth Position
Days
D
Average phase relative to neighbours
Average phase relative to neighbours
E
Local phase asymmetry
Local phase asymmetry
F
